# Reconciling simulated ensembles of apomyoglobin with experimental HDX data using Bayesian inference and multi-ensemble Markov State Models

**DOI:** 10.1101/563320

**Authors:** Hongbin Wan, Yunhui Ge, Asghar Razavi, Vincent A. Voelz

## Abstract

Hydrogen/deuterium exchange (HDX) is a powerful technique to investigate protein conformational dynamics at amino acid resolution. Because HDX provides a measurement of solvent exposure of backbone hydrogens, ensemble-averaged over potentially slow kinetic processes, it has been challenging to use HDX protection factors to refine structural ensembles obtained from molecular dynamics simulations. This entails two dual challenges: (1) identifying structural observables that best correlate with backbone amide protection from exchange, and (2) restraining these observables in molecular simulations to model ensembles consistent with experimental measurements. Here, we make significant progress on both fronts. First, we describe an improved predictor of HDX protection factors from structural observables in simulated ensembles, parameterized from ultra-long molecular dynamics simulation trajectory data, with a Bayesian inference approach used to retain the full posterior distribution of model parameters.

We next present a new method for obtaining simulated ensembles in agreement with experimental HDX protection factors, in which molecular simulations are performed at various temperatures and restraint biases, and used to construct multi-ensemble Markov State Models (MSMs). Finally, the BICePs algorithm (Bayesian Inference of Conformational Populations) is then used with our HDX protection factor predictor to infer which thermodynamic ensemble agrees best with experiment, and estimate populations of each conformational state in the MSM. To illustrate the approach, we use a combination of HDX protection factor restraints and chemical shift restraints to model the conformational ensemble of apomyoglobin at pH 6. The resulting ensemble agrees well with experiment, and gives insight into the all-atom structure of disordered helices F and H in the absence of heme.

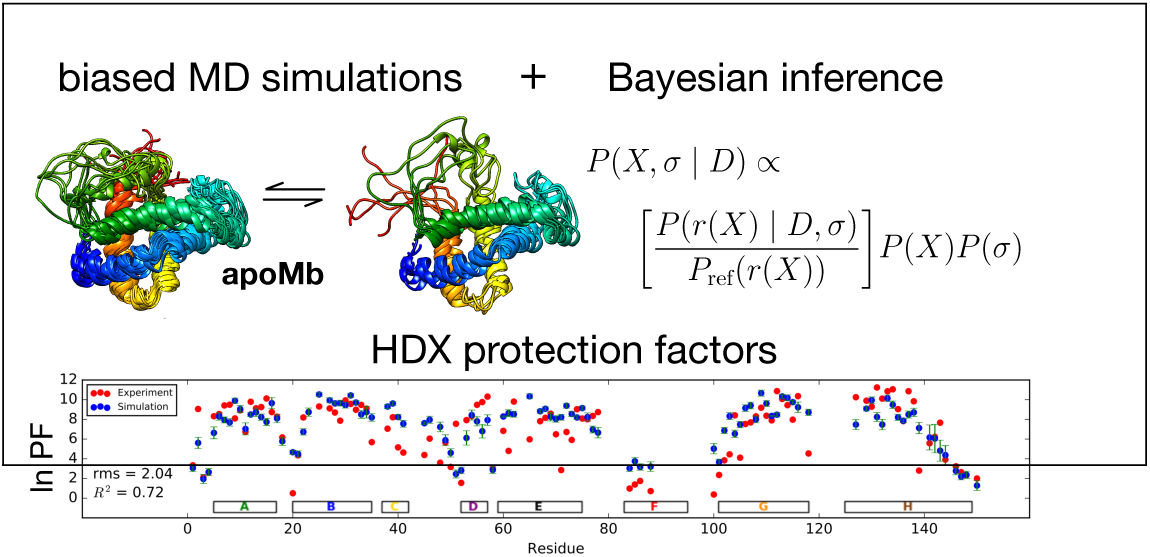
Graphical TOC Entry.

## Introduction

Hydrogen/deuterium exchange (HDX) is a powerful technique to investigate protein conformational dynamics at amino acid resolution. ^1–6^ In this technique, competition between the rates of exchange and the rates at which proteins exposing backbone amides can be used to probe a wide range of time scales, including very slow local and/or global unfolding/refolding dynamics. Exposed (unprotected) backbone amide hydrogens exchange with deuterated solvent according to the following kinetic model:

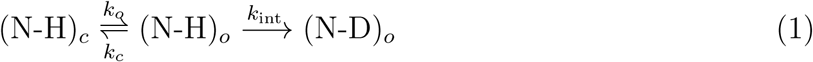

where *k*_*o*_ is the opening rate, *k*_*c*_ is the closing rate and *k*_int_ is the intrinsic exchange rate. The observed exchange rate, *k*_ex_ = *k*_*o*_*k*_int_*/*(*k*_*o*_ + *k*_*c*_ + *k*_int_), is very sensitive to temperature, pH, and the neighbor-dependent folded-state stability of each amino acid. ^7^

In the so-called EX1 regime, which occurs at high pH, high temperature or low stability, *k*_*c*_ *<< k*_int_, resulting in an exchange rate of *k*_ex_ = *k*_*o*_*k*_int_*/*(*k*_*o*_ + *k*_int_). In the so-called EX2 regime, *k*_*c*_ *>> k*_int_, i.e. the rate at which backbone amide hydrogens exchange with deuterium is slow compared to the rates at which backbone residues convert between “closed” conformations protected from exchange, and “open” conformations where exchange can occur. Therefore, the observed hydrogen/deuterium (HD) exchange rate, *k*_ex_ = *k*_*o*_*k*_int_*/*(*k*_*c*_ + *k*_int_), can be used to measure the relative populations of the “open” and “closed” states, by comparing it to the intrinsic exchange rate, *k*_int_, observed for an unstructured peptide. In this regime, the extent of protection for a residue *i* is characterized by a *protection factor*, 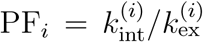, which can be related to the apparent free energy difference between open and closed states, Δ*G*, through ln PF_*i*_ = *β*Δ*G*_*i*_, where *β* = 1*/kT*.

## Existing methods for modeling HDX protection factors

Because HDX protection factors reflect potentially fleeting excursions to solvent-exposed states (“open” states), it has been challenging to make direct connections between molecular simulations of native-state protein dynamics and HDX protection factors, both in (1) predicting HDX protection factors directly from simulated trajectory data, and in (2) using experimental protection factors as restraints in simulated ensembles. Below, we review some of the methodology that has been used previously.

### Predicting HDX protection factors from trajectory data

Because most molecular simulations are unable to sample rare fluctuations on long time scales, much of the past work on predicting protection factors from trajectory data has relied on the correlation between structural observables and rare fluctuations, using proxy quantities such as solvent exposure and protein/solvent hydrogen bonding. Petruk et al. predicted protection from all-atom MD simulations of MAPK ERK2 protein using average solvent-accessible surface area and numbers of solvation waters for each backbone amide hydrogen as proxy structural variables. ^8^ Ma et al. have identified aggregation states of polymorphic amyloid *β*42 peptide through a combination of NMR HDX data and predicted protection factors using the ratio of the average number of hydrogen-bonds between amide hydrogens and water oxygens, and between amide hydrogens and carbonyl oxygens.^9^ Sljoka et al. use average hydrogen bond strengths to quantify protein rigidity/flexibility, which they use with solvent accessibility of backbone amide hydrogens to predict HDX data of Sso AcP from NMR ensembles.^10^ Kieseritzky et al. predict protection factors from MD simulations using hydrogen-bond occupancy, survival times, and fluctuations of backbone atoms and hydrogen bond length.^11^ Resing et al. showed that a linear combination of surface distance, inverse number of hydrogen-bonds, and the shortest distance to the first turn of the helix could predict the protection factors of ERK2 kinase helices with a linear correlation coefficient of 0.78.^12^

In a similar strategy, first employed by Vendruscolo et al., ^13^ experimental protection factors are modeled according to ln PF_*i*_ = *β*_*c*_⟨*N*_*c*_⟩_*i*_ + *β*_*h*_⟨*N*_*h*_⟩_*i*_, where ⟨*N*_*c*_⟩ _*i*_ is the average number of heavy-atom contacts with residue *i* and ⟨*N*_*h*_⟩_*i*_ is the average number of backbone hydrogen bonds. The parameters *β*_*c*_ and *β*_*h*_ can be determined by fitting the results of native-state protein simulations to experimental data.^13,14^ An advantage of this model is the computation of structural observables solely through pairwise distances, which are easily amenable to restraints. Another benefit of this model is its physical interpretation; the terms *β*_*c*_⟨*N*_*c*_⟩ _*i*_ and *β*_*h*_⟨*N*_*h*_⟩_*i*_ represent free energies of residue burial and hydrogen bonding, respectively.

Now that millisecond-long explicit-solvent MD trajectories have become available, ^15,16^ it has become possible to predict protection factors using a more mechanistic approach. Persson and Halle, ^17^ based on an analysis of the millisecond simulation trajectory of bovine pancreatic trypsin inhibitor (BPTI), have proposed that exchange-competent (“open”) conformations can be modeled as having two water oxygens found simultaneously within 2.6*Å* of the amide hydrogen. Persuasively, they show that direct counts of the number of trajectory snapshots containing open versus closed states gives computed protection factors in very good agreement with experiment. Limiting the practicality of the approach are (1) the need to obtain ultra-long simulation trajectories including sampled water configurations, and (2) the fact that highly protected amide hydrogens are likely coupled to global unfolding events which are not necessarily sampled in millisecond trajectories (indeed, such highly protected hydrogen exchange rates are not considered in the Persson and Halles analysis). It is problematic to use this “two-water” criterion to restrain simulated ensembles, as it would require three-body terms impractical for most molecular simulations. Nevertheless, the success of this approach suggests that millisecond simulations should provide more information for parameterizing empirical models than previously possible.

### Using experimental protection factors as restraints in simulated ensembles

Here, too, the inability of most simulations to sample fluctuations on long timescales makes it difficult to restrain ensemble-averaged structural observables correlated with backbone amide protection. One approach has been to use simple structural models enabling the enumeration of a complete statistical thermodynamic ensemble. In the DXCOREX method of Liu et al,^18^ the statistical thermodynamic ensemble of the protein is modeled as a set of folding units (microstates) that are either folded or unfolded, allowing complete enumeration of the complete state space and state probabilities according to an empirical Gibbs free energy function that depends on the accessible surface area of polar and nonpolar microstates. The per-residue protection factors can then be calculated from the Boltzmann probabilities of folded vs. unfolded states.

Another approach is to use restraint-biased all-atom simulations to model structural ensembles. Typically, these methods are used to achieve partial or global unfolding of a protein to produce ensembles more consistent with experimental protection factors. To restrain ensemble-average quantities in all-atom molecular dynamics ensembles, Vendruscolo et al. ^13^ developed a method whereby multiple simulation replicas are simultaneously maintained, with harmonic restraints enforcing the average ⟨ln PF_*i*_ ⟩= *β*_*c*_⟨*N*_*c*_⟩_*i*_ +*β*_*h*_⟨*N*_*h*_⟩_*i*_ calculated across all the simulation replicas. All-atom simulations of chymotrypsin inhibitor 2 (CI2) restrained by this method yield conformational ensembles consistent with experiment.

One problem with restraint simulations is the risk of introducing unnecessary bias into the ensemble from the restraint potential. Pitera and Chodera have used a maximum entropy approach to show that the least-biased method to restrain some ensemble-averaged quantity ⟨*f* (*x*) ⟩, where *f* (*x*) is a structural observable computed for a conformation, *x*, is to use a modified force field potential *U* (*x*) = *U* (*x*) + *αf* (*x*), for some scaling parameter *α*.^19^ In practice, the value of *α* can be determined by performing multiple simulations at different values of *α*, and selecting the value that reproduces the correct value of ⟨*f* (*x*) ⟩. In the limit of large numbers of replicas, the Vendroscolo et al. method approaches this maximum entropy solution. The maximum entropy method has a practical drawback, however: using it to restrain protection factors for a large number of amino acids in a protein would require exploring an enormously large parameter space. As we show below, we can alternatively use a simplified version of this idea to simulate ensembles more consistent experimental protection factors.

## Overview

In this manuscript, we expand on previous work in several ways. Our results are organized into three parts. In Part I, we take a Bayesian inference approach to parameterizing an empirical predictor of HDX protection factors from molecular simulation data. Starting with a functional form similar to Vendruscolo et al., we fit against newly-available ultralong molecular simulation trajectory data, while retaining the full posterior distribution of model parameters. In Part II, we pursue a new way of performing biased simulations to generate structural ensembles consistent with HDX protection factor data, through the example of apomyoglobin (apoMb). Inspired by the maximum entropy method of Pitera and Chodera,^19^ we perform simulations of apoMb using a number of different bias potentials and temperatures, and use the resulting trajectory data to construct multi-ensemble Markov State Models.^20^ In Part III, we use a Bayesian inference approach, implemented through our BICePs (Bayesian Inference of Conformational Populations) algorithm, ^21,22^ to reconcile the MSMs built for each thermodynamic ensemble against experimental protection factor measurements and chemical shift measurements. The key advantage of this approach is that we can use Bayesian inference to propagate uncertainty in model parameters (found in the first part) to perform quantitative model selection.

## Methods and Results

### Part I: An empirical model of HDX protection parameterized from ultra-long simulation trajectory data

We first attempted to construct a new empirical model–trained on ultra-long MD trajectories– to predict protection factors according to the following form:

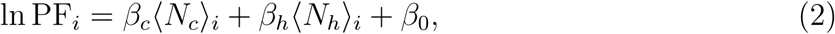

This form is similar to the Vendruscolo et al. model, ^13^ but with an additional cooperativity term *β*_0_ that can compensate for the correlations between heavy-atom contacts and hydrogenbond contacts.

The values of the parameters for this model come from fitting to ultra-long (millisecond) native-state molecular dynamics simulations of the 58-residue protein BPTI,^15^ and the 76residue protein ubiquitin,^16^ both provided by D.E. Shaw Research. First, we will describe the simulation trajectory data sets, and later describe our parameterization scheme, in which we use Bayesian inference to compute the full posterior distribution of likely parameters.

#### BPTI and ubiquitin molecular dynamics trajectory data

From each simulation, we procured a sample trajectory of 50000 snapshots for model parameterization, typical of conventional explicit-solvent simulation trajectories. The native-state BPTI simulation was performed at 300 K with 4215 water molecules, from which we analyzed a segment of the full trajectory (71-83.5 *µ*s) containing 50000 frames taken every 250 ps. The native-state ubiquitin simulation was performed at 300 K with 5581 water molecules, from which we analyzed a trajectory of 50000 snapshots taken every 20 ns. The RMSD variances on the native state ensembles of both systems is small (Figure 1), and thus can be used to study hydrogen exchange in native states.

**Figure 1:**
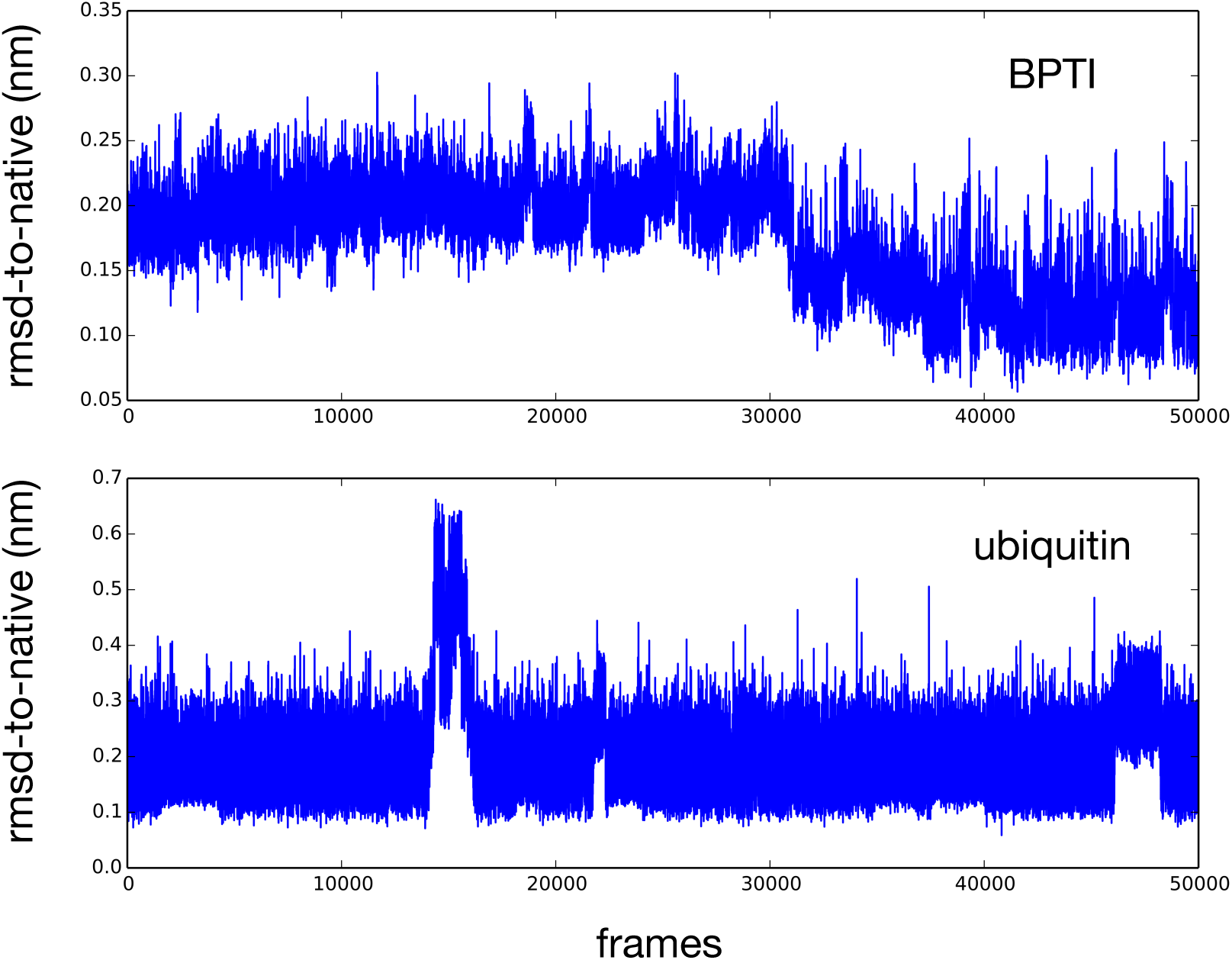
Molecular dynamics trajectory data used for training an empirical model of HDX protection. Shown is rmsd-to-native over time for 50000 frames of BPTI (frames every 250 ps) ubiquitin (frames every 20 ns). Trajectory data was provided by D.E. Shaw Research.

#### Bayesian model parameterization

In addition to the scaling parameters *β*_*c*_, *β*_*h*_, and *β*_0_ in Equation (2), there are parameters associated with how the average numbers of heavyatom contacts ⟨*N*_*c*_⟩_*i*_ and hydrogen-bonds ⟨*N*_*h*_⟩_*i*_ are computed for each residue *i*. The averages are calculated using sigmoidal cut-off functions averaged over all *T* snapshots in the sample. ⟨*N*_*c*_⟩_*i*_ is computed as

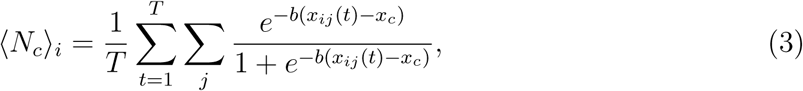

where *x*_*ij*_(*t*) are the distances (in Å) from the backbone amide nitrogen of residue *i* to other heavy atoms *j*, at snapshot *t*, and *x*_*c*_ is a distance threshold parameter defining a heavy-atom contact. Similarly, ⟨ *N*_*h*_⟩ _*i*_ is computed as:

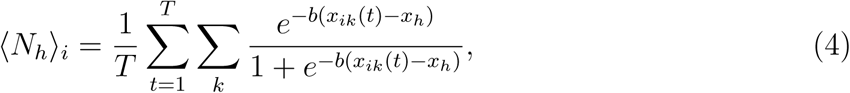

where *x*_*ik*_(*t*) are the distances (in Å) between the backbone amide hydrogen of residue *i* to oxygen hydrogen-bond acceptors *k*, at snapshot *t*, and *x*_*h*_ is a distance threshold parameter defining a hydrogen bond. The parameter b has units Å^*-*1^ and controls the sharpness of the sigmoidal cutoff for both ⟨ *N*_*ci*_ ⟩and ⟨*N*_*h*_⟩_*i*_. Taken together, there are six parameters in our model to be inferred from the training data: *β*_*c*_, *β*_*h*_, *β*_0_, *x*_*c*_, *x*_*h*_, and *b*.

To determine these parameters, we implement a Bayesian inference approach. While traditional optimization schemes aim to find a particular set of parameters that maximize a likelihood function, Bayesian approaches aim to sample the entire posterior distribution of parameters *λ* = (*β*_*c*_, *β*_*h*_, *β*_0_, *x*_*c*_, *x*_*h*_, *b*), from which uncertainty estimates can be computed. By Bayes theorem, the posterior probability distribution *P* (*λ|D*) of parameters, given some experimental data *D*, obeys the proportionality

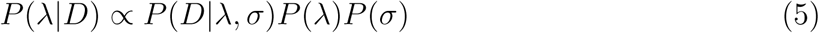

where *P* (*D|λ*) is a likelihood function describing the probability of observing the data given the parameters, and *P* (*λ*) is a prior distribution of parameters, which we set to be uniform in some reasonable range. For our likelihood function, we use a Gaussian error function

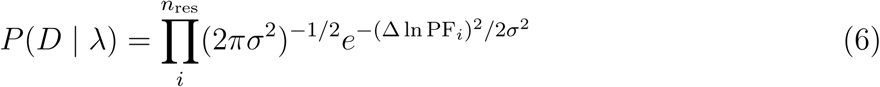

where Δ(ln PF_*i*_) are the differences in experimental and predicted protection factors, *n*_res_ is the number of residues, and *σ* is a parameter specifying the expected error. Since the expected error is unknown, we include *σ* as a nuisance parameter in the posterior distribution,

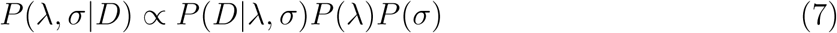

with *P* (*σ*) *∼ σ*^*−*1^ chosen to be an uninformative Jeffreys prior.

#### Training the model

The experimental data used to train the model included: (1) 72 experimental protection factors for ubiquitin compiled by Craig et al. as the average of rescaled HDX data studied at different pH,^23^ and (2) experimental protection factors for 30 of the 53 amide hydrogens of BPTI with published NMR HDX measurements at 300 K.^24–27^ These 30 were the same set used by Persson and Halle, in which the authors excluded highly protected amides and surface amides which exhibited anomalous pH dependence. ^17^ A full list of experimental protection factors, converted to ln PF values, are listed in the Supporting Information (Tables S1 and S2).

Training the model entails sampling the full posterior *P* (*λ, σ|D*) over all model parameters (*λ* = (*β*_*c*_, *β*_*h*_, *β*_0_, *x*_*c*_, *x*_*h*_, *b*), *σ*) using random walk Monte Carlo sampling. At each step, one of these seven variables was randomly chosen and a move was proposed to a new nearestneighbor on a grid of allowed values, and accepted with the Metropolis criterion. Values of *β*_*c*_ ranged from 0.05 to 0.20 *kT* in increments of 0.01 *kT*. Values of *β*_*h*_ ranged from 0 to 5.0 *kT* in increments of 0.2 *kT*. Values of *β*_0_ ranged from −10 to 0 *kT* in increments of 0.2 *kT*. Values of *x*_*c*_ ranged from 5.0 to 8.0 Å in increments of 0.5 Å. Values of *x*_*h*_ ranged from 2.0 to 2.7 Å in increments of 0.1 Å. Values of *b* ranged from 3 to 20 Å^*-*1^ in increments of 1 Å^*-*1^. The value of *σ* was constrained to 100 log-spaced grid values from 0.25 to 5.0. Using these values, all variables had acceptance ratios greater than 0.50. Trials of 10^6^ and 10^7^ steps were performed, with similar results (see below).

The marginal distribution of *P* (*λ|D*) is obtained through ∫*P* (*λ, σ|D*)*dσ*. Posterior marginal distributions for each parameter are calculated similarly, from the values sampled by Monte Carlo (Figure 2). To ensure robust results, we trained the model separately on ubiquitin (Figure 2a) and BPTI (Figure 2b), and also on both data sets (Figure 2c).

**Figure 2:**
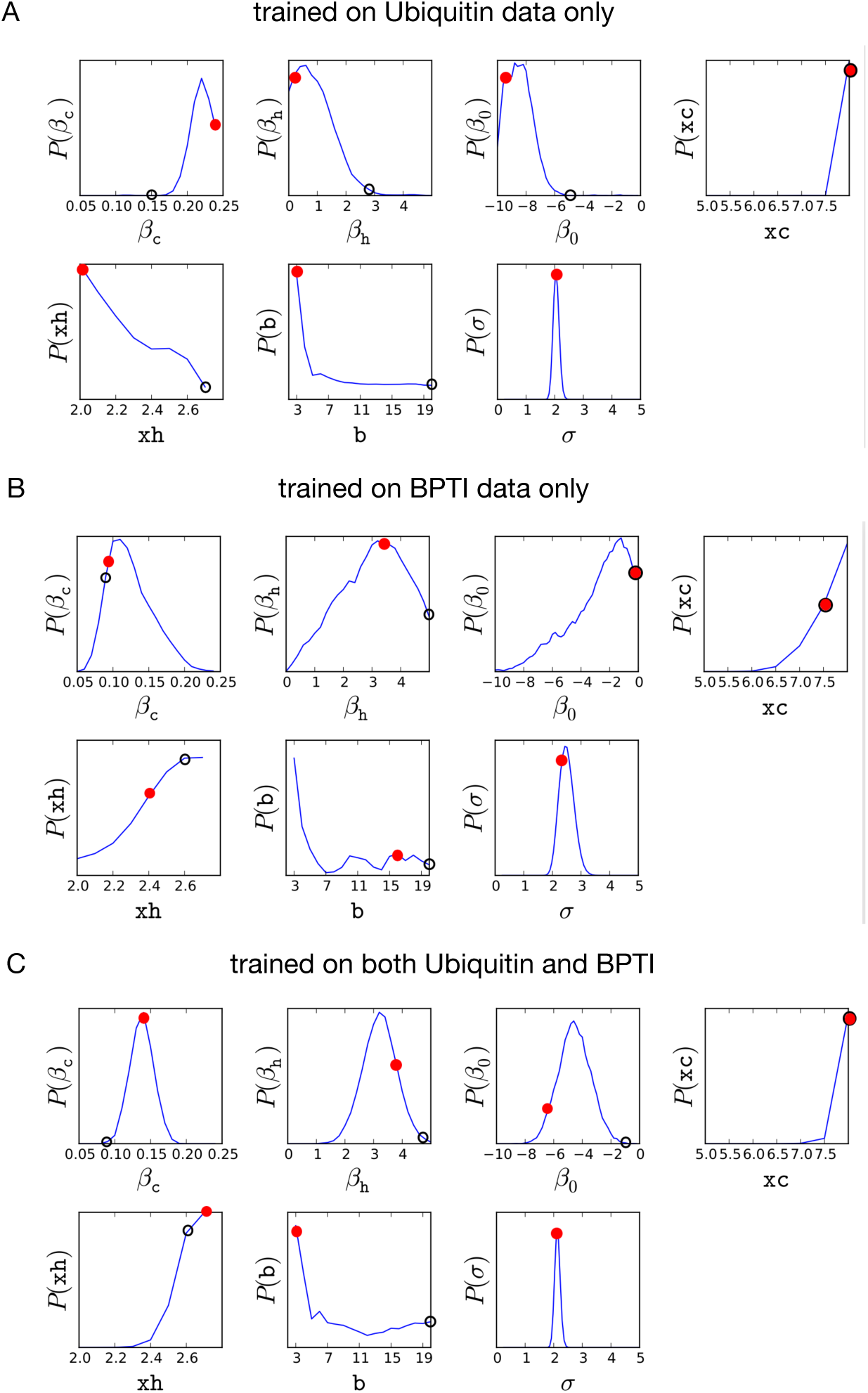
Posterior marginal distributions of model parameters *β*_*c*_*β*_*h*_, *β*_0_, *x*_*c*_, *x*_*h*_, *b*, and *σ*, obtained from Monte Carlo sampling. Parameters *β*_*c*_*β*_*h*_, *β*_0_ are in units *kT*. Parameters *x*_*c*_ and *x*_*h*_ have units Å. The parameter *b* (units Å^*-*1^) controls the sharpness of the sigmoidal cutoff for determining the presence of a heavy-atom contact or hydrogen bond, and *σ* represents the standard error in predicting ln PF_*i*_.

The posterior distributions of model parameters are similar when trained individually on ubiquitin or BPTI protection factors alone, but differences can be observed, mainly in the importance of heavy-atom coordination versus hydrogren bonds. Whereas the ubiquitin trained model has a large *β*_*c*_ coefficient and a low *β*_*h*_ coefficient (with a lower distance threshold *x*_*h*_ for including hydrogen bonds), the opposite is true for the BPTI-trained model. The model trained on both data sets has posterior distributions centers on intermediate values of *β*_*c*_ and *β*_*h*_.

Of all the possible sets of parameters sampled in the full posterior distribution, it is useful to pick a single set of parameters to formulate a ln PF_*i*_ predictor. We do this by choosing the maximum a posteriori (MAP) parameter values *λ*^*\**^ = argmax *P* (*λ|D*), i.e. the parameters that give the maximum value of the joint posterior distribution. The MAP is distinguished from the maximum likelihood (ML) parameters, which are the set of parameters that minimize the likelihood function in Equation 6. Because this likelihood function is a Gaussian error function, it is minimized when sum of squared errors (SSE) Σ*i* Δ ln PF_*i*_ is minimized. Thus, the ML model is comparable to a linear regression model where the sum of squared residuals are minimized. Such models can be overly sensitive to outliers, a problem which can be ameliorated with the use of a Bayesian posterior. Indeed, we find that the ML model (black circles in Figure 2) parameters are located in the tails of the posterior distribution, unrepresentative of the larger posterior distribution. Moreover, we also note that, because the parameters are not independent, the maximum a posteriori (MAP) set of parameters *λ*^*\**^ = argmax *P* (*λ|D*) (red filled circles in Figure 2) is *not* the maximum of each marginal posterior distribution.

ML and MAP parameters *λ*^*\**^ for ubiquitin-trained (Ubq) and BPTI-trained models (BPTI) are shown in Tables 1 and 2, respectively. ML and MAP parameters *λ*^*\**^ for models trained on both data sets (Ubq+BPTI) are shown in Table 3. To test whether we have adequately sampled the posterior distribution, we compare the results when using 10^6^ and 10^7^ MCMC samples; the results are extremely similar in all cases. The rms errors in the ln PF_*i*_ predictions from MAP are not that much larger than those for the ML models (which by definition give the lowest rms errors): whereas the Ubq, BPTI and (Ubq+BPTI) ML models yield rms errors in ln PF_*i*_ of 2.555, 2.714 and 2.737, respectively, the MAP models have rms only slightly larger: 2.766, 3.011 and 2.914, respectively (using 10^7^ MCMC samples). For both the ML and MAP models, training on both sets of data (Ubq+BPTI) yields rms errors similar to models trained on each protein alone (Ubq, or BPTI).

**Table 1:**
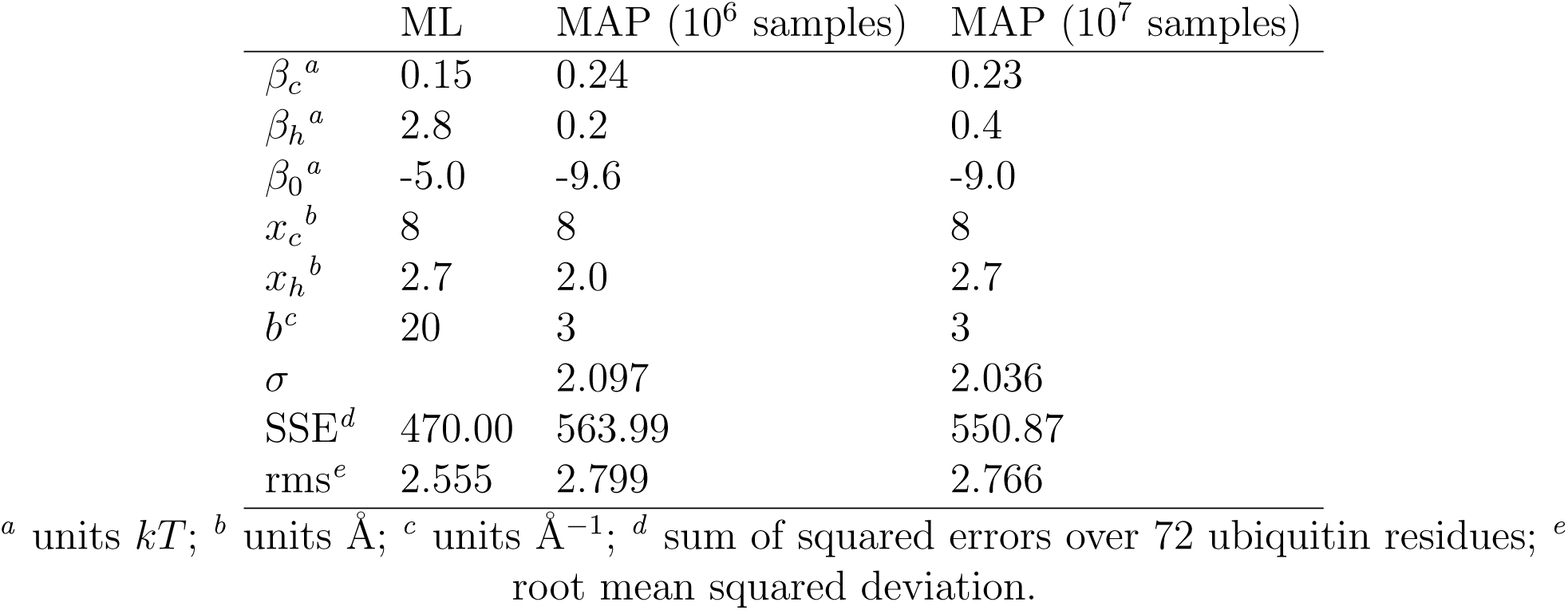
Maximum likelihood (ML) and maximum a posteriori (MAP) model parameters for a ln PF_*i*_ predictor trained on ubiquitin data only (Ubq).

| | ML | MAP ($10^6$ samples) | MAP ($10^7$ samples) |
| --- | --- | --- | --- |
| $\beta_c^a$ | 0.15 | 0.24 | 0.23 |
| $\beta_h^a$ | 2.8 | 0.2 | 0.4 |
| $\beta_0^a$ | -5.0 | -9.6 | -9.0 |
| $x_c^b$ | 8 | 8 | 8 |
| $x_h^b$ | 2.7 | 2.0 | 2.7 |
| $b^c$ | 20 | 3 | 3 |
| $\sigma$ | | 2.097 | 2.036 |
| SSE<sup>d</sup> | 470.00 | 563.99 | 550.87 |
| rms<sup>e</sup> | 2.555 | 2.799 | 2.766 |

<sup>a</sup> units $kT$ ; <sup>b</sup> units $\text{\AA}$ ; <sup>c</sup> units $\text{\AA}^{-1}$ ; <sup>d</sup> sum of squared errors over 72 ubiquitin residues; <sup>e</sup> root mean squared deviation.

**Table 2:** Maximum likelihood (ML) and maximum a posteriori (MAP) model parameters for a ln PF_*i*_ predictor trained on BPTI data only (BPTI).

| | ML | MAP ($10^6$ samples) | MAP ($10^7$ samples) |
| --- | --- | --- | --- |
| $\beta_c^a$ | 0.08 | 0.09 | 0.07 |
| $\beta_h^a$ | 5.0 | 3.4 | 4.0 |
| $\beta_0^a$ | -0.2 | -0.2 | -1.8 |
| $x_c^b$ | 7.5 | 7.5 | 8.0 |
| $x_h^b$ | 2.6 | 2.4 | 2.7 |
| $b^c$ | 20 | 16 | 3 |
| $\sigma$ | | 2.227 | 2.227 |
| SSE<sup>d</sup> | 221.03 | 267.58 | 272.00 |
| rms<sup>e</sup> | 2.714 | 2.986 | 3.011 |

<sup>a</sup> units $kT$ ; <sup>b</sup> units $\text{\AA}$ ; <sup>c</sup> units $\text{\AA}^{-1}$ ; <sup>d</sup> sum of squared errors over 30 BPTI residues; <sup>e</sup> root mean squared deviation.

**Table 3:** Maximum likelihood (ML) and maximum a posteriori (MAP) model parameters for a ln PF_*i*_ predictor trained on both ubiquitin and BPTI data (Ubq+BPTI).

| | ML | MAP ($10^6$ samples) | MAP ($10^7$ samples) |
| --- | --- | --- | --- |
| $\beta_c^a$ | 0.08 | 0.14 | 0.14 |
| $\beta_h^a$ | 4.6 | 3.8 | 3.4 |
| $\beta_0^a$ | -1 | -6.2 | -5.8 |
| $x_c^b$ | 8.0 | 8.0 | 8.0 |
| $x_h^b$ | 2.6 | 2.7 | 2.7 |
| $b^c$ | 20 | 3 | 3 |
| $\sigma$ | | 2.161 | 2.161 |
| SSE<sup>a</sup> | 764.25 | 872.10 | 866.14 |
| rms<sup>b</sup> | 2.737 | 2.924 | 2.914 |

<sup>a</sup> units $kT$ ; <sup>b</sup> units $\text{\AA}$ ; <sup>c</sup> units $\text{\AA}^{-1}$ ; <sup>d</sup> sum of squared errors over 102 residues; <sup>e</sup> root mean squared deviation.

When we use the MAP *λ*^*\**^ of Ubq-, BPTIand (Ubq+BPTI)-trained models to compare experimental and predicted values of ln PF_*i*_ (Figure 3), we find squared correlation coefficient *R*^2^ values of 0.312, 0.480 and 0.466, respectively. Since the (Ubq+BPTI)-trained MAP model exhibits the best balance of low rms error and high *R*^2^ values, and uses all the available training data, we use this model for all subsequent work (see Part III). The *R*^2^ value of this model is comparable to Craig et al.’s prediction on ubiquitin (*R*^2^ = 0.53) and Persson and Halle’s predictions for BPTI (*R*^2^ = 0.68). The lower extent of correlation for our PF predictions may arise in part from our smaller training set of molecular simulation data. Only 5 × 10^4^ frames were used to train our model, versus 10^6^ frames used in Halle’s predictions on BPTI, and 2 × 10^5^ frames for Craig’s prediction on ubiquitin.

**Figure 3:**
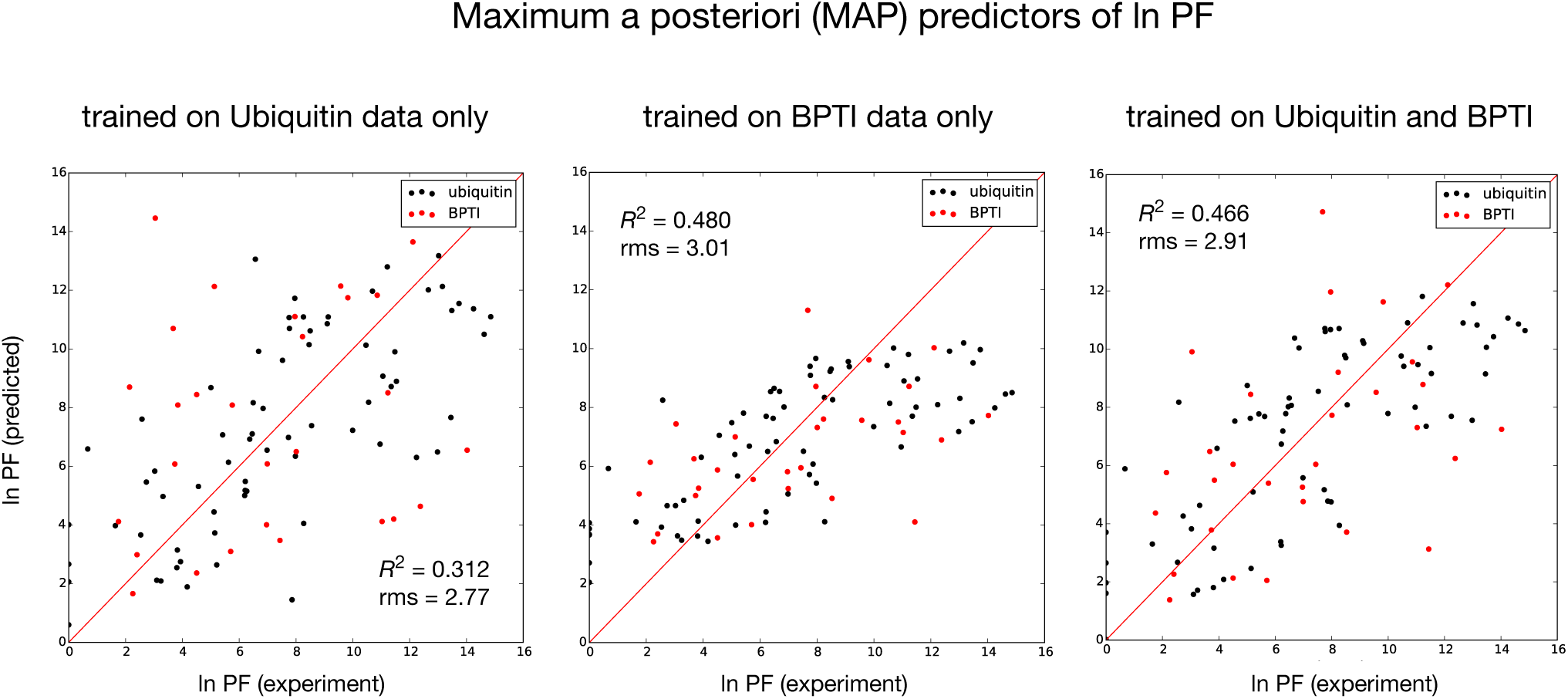
Comparisons of experimental and predicted values of ln PF_*i*_ from maximum a posteriori (MAP) models trained on ubiquitin data only (Ubq), BPTI data only (BPTI), and both data sets (Ubq+BPTI). Values of *R*^2^ and rms values reported in each subplot are for the entire dataset (Ubq+BPTI)

The values of *β*_*c*_ and *β*_*h*_ in our final MAP (Ubq+BPTI) model (Table 3, 10^7^ steps) can be used to gain insight into the unfolding (closed-to-open state) free energy contributions provided by heavy-atom contacts and hydrogen bonds, through the terms *β*_*c*_ *·* ⟨*N*_*c*_⟩_*i*_ and *β*_*h*_ *·* ⟨*N*_*h*_⟩_*i*_, respectively, for each residue *i* (Figure 4). These values range from contributions of 4–16 *kT* attributed to heavy-atom contacts, and 1–6 *kT* attributed to hydrogen bond breaking, depending on the residue. These two free energy contributions are correlated, as indicated by the cooperativity term of *β*_0_ = −5.8 *kT*. This term provides a correction factor to offset the “double-counting” of these related contributions. These results are similar to the results of Vendruscolo et al., who estimated (uncorrelated) free energy contributions of 0.6 kcal mol^*-*1^ (*∼*1 *kT*) per heavy-atom contact, and 3 kcal mol^*-*1^ (*∼*5 *kT*) per hydrogen bond.^13^

**Figure 4:**
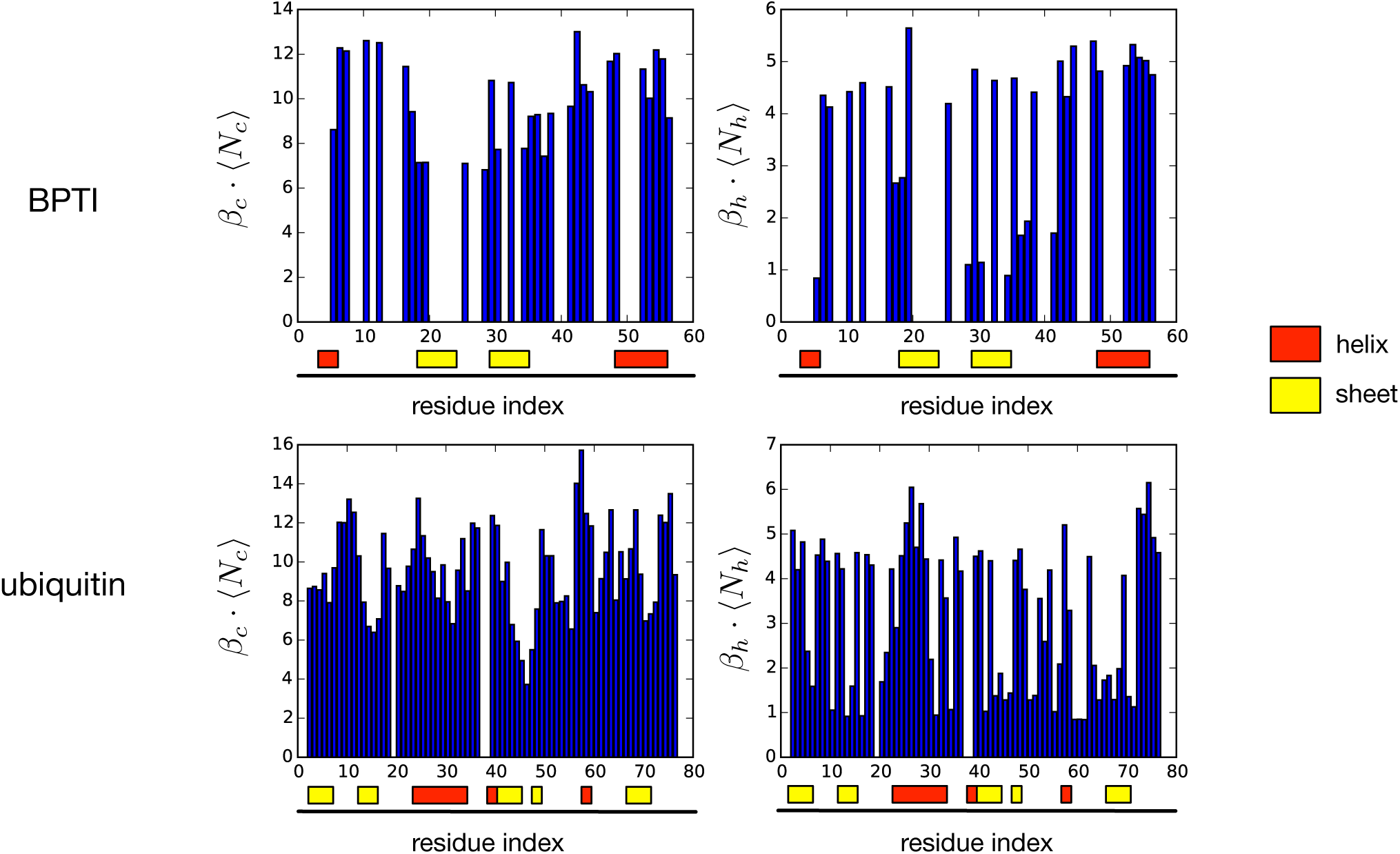
Estimates of the unfolding (closed-to-open state) free energy contributions (in units *kT*) originating from heavy-atom contacts *β*_*c*_ *·* ⟨*N*_*c*_⟩_*i*_, versus hydrogen bonding *β*_*h*_ *·* ⟨*N*_*h*_⟩_*i*_ for each residue *i*, according to the MAP predictor of ln PF_*i*_.

### Part II. Restraint-biased simulation and construction of multi-ensemble MSMs for apoMb

From our work above, we now have in hand a reasonably accurate function, ln PF_*i*_(*X*) = *β*_*c*_⟨*N*_*c*_⟩_*i*_(*X*)+*β*_*h*_⟨*N*_*h*_⟩_*i*_(*X*)+*β*_0_, that yields a prediction of ln PF_*i*_ for residue *i*, given a molecular conformation *X*. According to the maximum entropy approach of Pitera and Chodera, ^19^ the least-biased potential to restrain protection factor observables in a molecular simulation is expressed as a modified potential *U ′*(*X*) = *U* (*X*) +Σ*_i_ α*_*i*_(ln PF_*i*_(*X*)). This would require performing an unfeasible number of simulations to explore the full parameter space of all *α*_*i*_.

Instead, we propose a simplification to this scheme, in which a single restraint bias potential (with a single parameter *α*), is applied to multiple protein residues, so as to generate structural ensembles with different extents of solvent exposure and amide hydrogen bonding. Later (as we describe below), the ensembles will be evaluated using the BICePs algorithm to determine which is most consistent with the experimental data.

#### Apomyoglobin

As a specific system on which to test this approach, we consider apomyoglobin (apoMb), a protein whose folding has been well-studied by NMR and x-ray crystallography.^28,29^ Myoglobin is a 152-residue heme protein with eight helices labeled A through H (Figure 5). In the absence of heme at pH 6, apomyoglobin adopts a holoprotein-like conformation, although the F helix and C-terminal portion of the H helix becomes disordered.^30,31^ This conformation is known as the *native* (*N*) state of apoMb. At pH 4.0, apomyoglobin becomes more highly disordered; this acid-denatured state (M) is similar to a kinetic intermediate in the refolding of apoMb, as characterized by quench-flow amide proton H/D exchange pulse labeling and stopped-flow spectroscopy. ^32–36^

**Figure 5:**
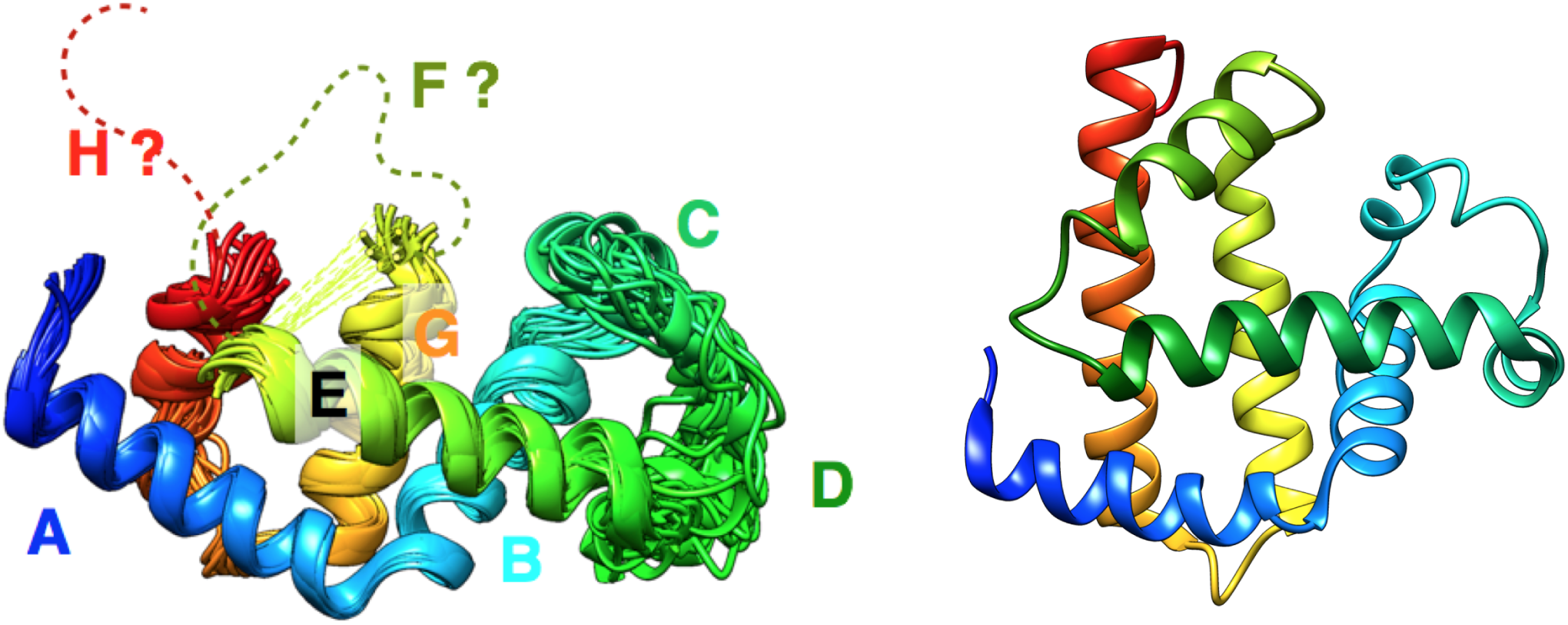
(Left) NMR structure of apomyoglobin at neutral pH shows disordered F and H helices (Lecomte 1999, personal communication). (Right) Holo-myoglobin (PDB:1JP6) was the starting conformation of the restraint-bias molecular simulations.

Here, we focus on generating simulated ensembles of apoMb that best represent the N state of apoMb. Our primary goal is to reconcile the ensembles against protection factors for apomyoglobin at pH 6.0 and pH 4.0 measured by Nishimura et al. ^37^ and NMR chemical shifts measured by Eliezer and Wright.^30^

#### Molecular simulation

Molecular simulations of apomyoglobin were prepared and performed using the GROMACS 5.0.1 simulation package on TACC Stampede supercomputer. NVT simulations were performed using a stochastic (Langevin) integrator with step size 2 fs. The AMBER ff99SB-ildn-nmr force field and TIP3P water model were used with cubic periodic box of volume (6.743 nm)^3^ containing 30072 atoms, which included the protein, 9194 water molecules, 18 Na^+^ ions, and 20 Cl^*-*^ ions (approximately 100 mM salt concentration).

The starting conformation of the protein was taken from holomyoglobin (PDB:1JP6). Protonation states at pH 7 were chosen according to the pKa values measured by Geierstanger et al. ^32^

#### Restraint bias potentials to encourage solvent exposure

To encourage the solventexposure of specific residues, sigmoidal restraint bias potentials were included in the simulations (Figure 6). The restraint biases were implemented using tabulated bonded interactions (cubic spline potentials) in GROMACS, of the form *U*_bias_(*x*) = *k · f* (*x*), where *k* is a force constant in units of energy, and *f* (*x*) is a function of interatomic distance *x*,

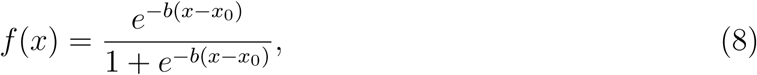

where *x*_0_ = 0.65 nm, and *b* = 5 nm^*-*1^.

**Figure 6:**
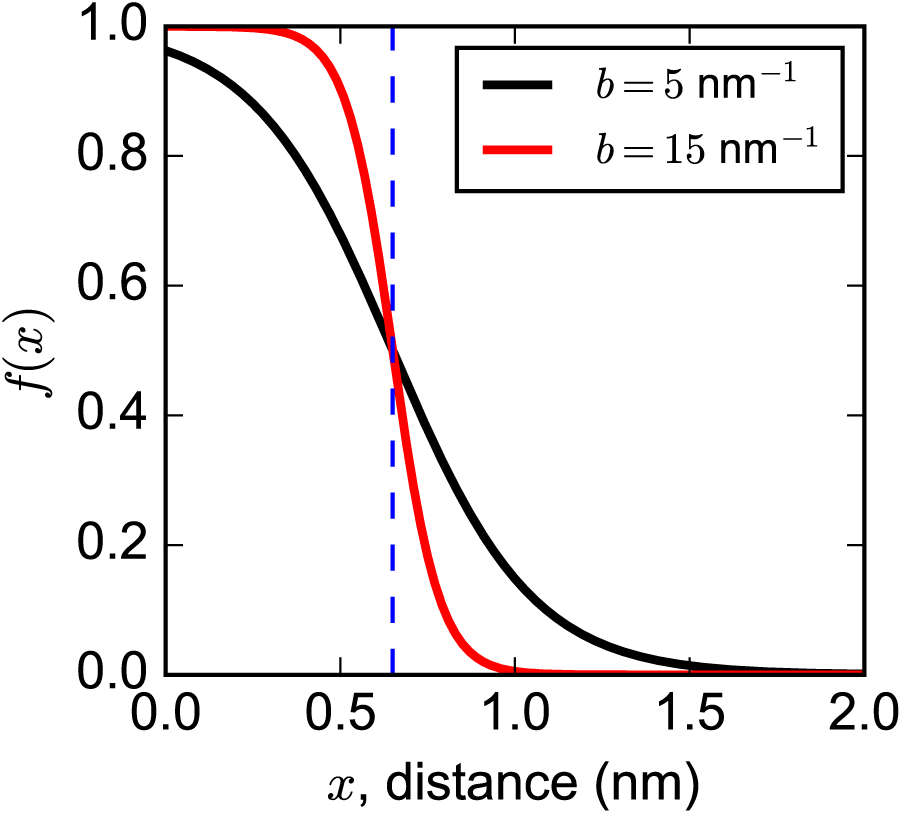
The sigmoidal function *f* (*x*) used with the biasing potential *U*_bias_(*x*) = *k · f* (*x*), where *k* is a force constant in units of energy. The value of *b* was set to 5 nm^*-*1^.

Protection factors for apoMb measured by Nishimura et al. ^37^ show that helix F and the C-terminal region of helix H are more solvent-exposed for apoMb than the *holo* protein. Therefore, bias restraints were added between the amide hydrogens of in helix F (residues 83-87, 89-95), and helix H (residues 140-152) and oxygens on all residues capable of making hydrogen bonds; these included backbone carbonyl oxygens, as well as side chain oxygen atoms on aspartic acid, glutamic acid, glutamine, serine, threonine and tyrosine (Figure 7).

**Figure 7:**
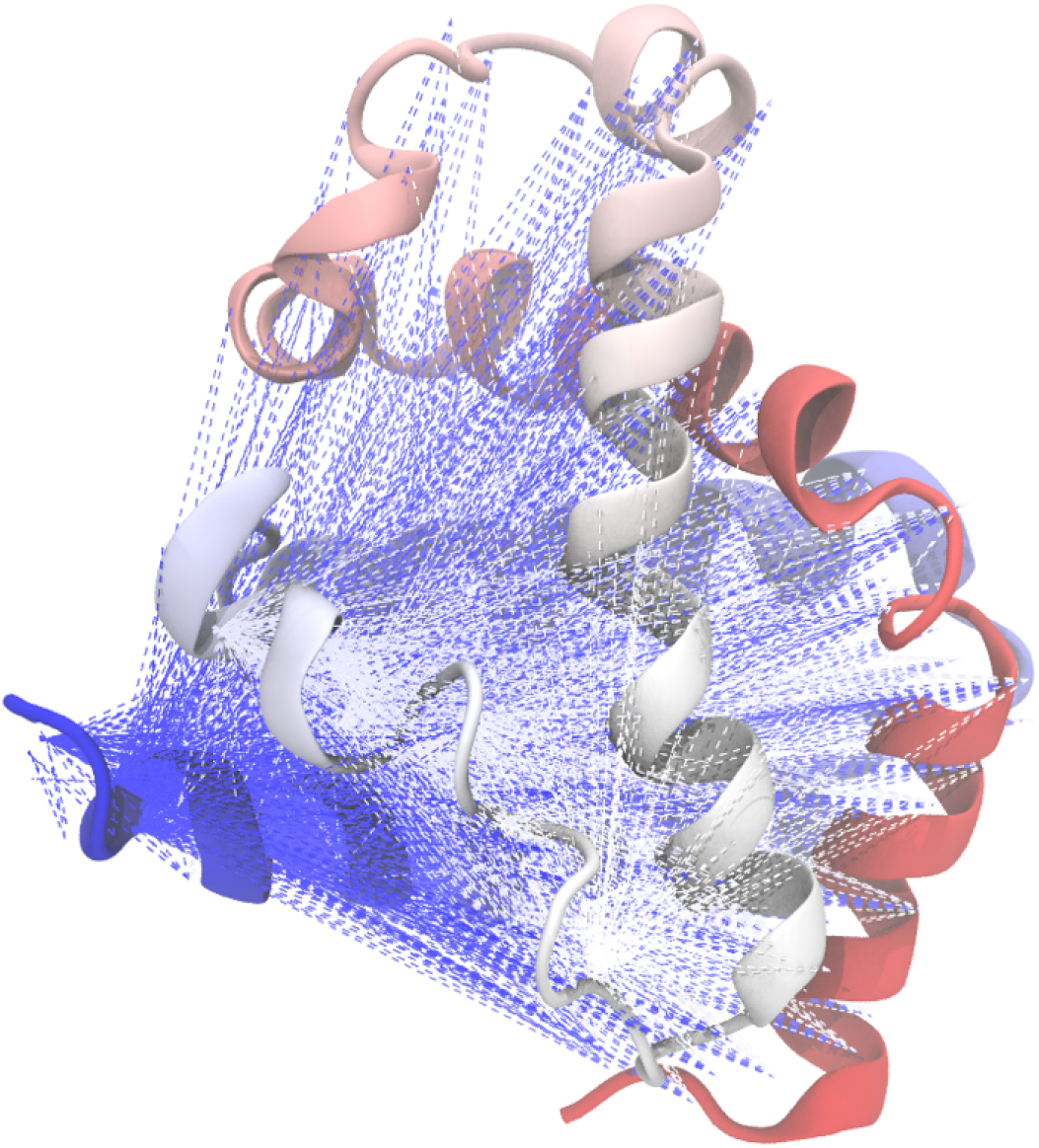
Visualization of protection factor restraints applied to selected residues of apoMb.

Simulations were performed at temperatures 300, 330, 350, 370, 400 and 415 K. For each temperature, simulations were performed using force constants of *k* = 0.5, 0.7, 1.0, 1.2, 1.5 and 2.0 kJ, resulting in 36 simulations totalling 19 *µ*s of aggregate simulation trajectory data (Table 4), with snapshots saved every 100 ps.

**Table 4:** A summary of molecular dynamics simulation trajectory data for apoMb.

| force constant (kJ) | trajectory length (in μs) | | | | | |
| --- | --- | --- | --- | --- | --- | --- |
|  | 300 K | 330 K | 350 K | 370 K | 400 K | 415 K |
| 0.0 | 15.7 | 6.2 | 5.9 | 5.6 | 6.2 | 6.0 |
| 0.5 | 1.0 | 1.0 | 1.0 | 0.8 | 0.08 | 0.08 |
| 0.7 | 1.0 | 1.0 | 1.0 | 0.8 | 0.08 | 0.08 |
| 1.0 | 1.0 | 1.0 | 1.0 | 0.07 | 0.08 | 0.08 |
| 1.2 | 1.0 | 1.0 | 1.0 | 0.7 | 0.08 | 0.08 |
| 1.5 | 1.0 | 1.0 | 1.0 | 0.08 | 0.08 | 0.08 |
| 2.0 | 0.6 | 1.0 | 1.0 | 0.08 | 0.08 | 0.08 |

### Construction of multi-ensemble MSMs from restraint-biased trajectory data

The simulation trajectory data obtained at multiple temperatures and restraint-bias potentials were next used to construct multi-ensemble Markov Models (MEMMs). These models can be thought of as a *set* of MSMs–one for each thermodynamic ensemble. The main advantage of MEMMs is that observed transitions sampled across all the ensembles can provide information to estimate transition rates between states in each individual ensemble.

#### Projection of trajectory data to discrete metastable states

Discretization of the trajectory data for MSM analysis was accomplished by first performing dimensionality reduction of the coordinate data, and then conformational clustering in the low-dimensional projection to define metastable states.

Time-lagged independent component analysis (tICA) ^38,39^ was used to determine the lowdimensional subspace corresponding to the slowest motions of the protein. Similar to principal component analysis (PCA), which finds the eigenvectors of a covariance matrix, the tICA method solves a similar eigenvalue problem for a time-lagged correlation matrix to find the degrees of freedom that capture the most time-correlated motions. As input coordinates, we used all pairwise distances between C_*α*_ atoms. The tICA lag time used was 0.5 ns. The entire set of trajectory data (all temperatures and force constants) was used as input to tICA.

Next, the trajectory data were projected to the top 8 tICA components, and *k*-centers clustering was performed in this subspace to identify 25 microstates with which MSMs (and MEMMs) could be constructed (Figure 8). The number of microstates was chosen to facilitate sufficient overlap of metastable states to construct MEMMs (as described below). Visualization of the trajectory data on tIC_1_ and tIC_2_ shows that the slowest motion (moving left to right along tIC_1_) corresponds to the unstructuring of helix F and helix H. Microstate 16 (microstates are numbered using indices 0 through 24) on the left side of Figure 8a is the state from which all trajectories were initiated, a *holo*-like structure with a folded F-helix and an rmsd of 0.25 nm to the native structure. On the right side of Figure 8a are a number of microstates corresponding to conformational states lacking structure in helix F and H.

**Figure 8:**
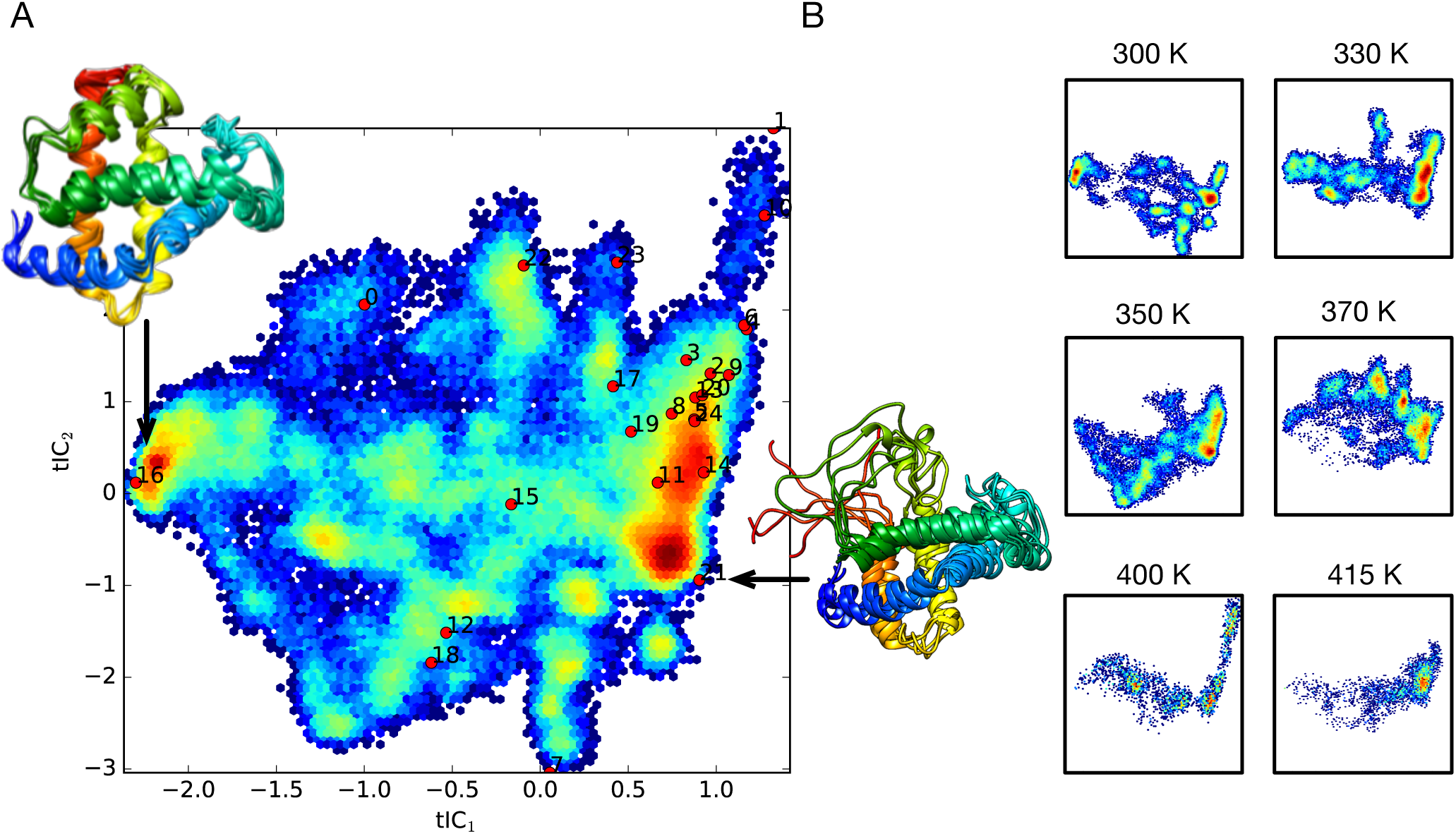
Visualization of apoMb trajectory data on the tICA landscape. (A) A density map of trajectory data from all temperatures and force constants projected to tIC_1_ and tIC_2_. Red dots indicate the centers of the conformational clusters used as MSM metastable states, labeled by microstate index. (B) Density maps of trajectory data for the six simulation temperatures, shown separately on the same axes as panel A.

#### Construction of MEMMs

The TRAM (transition-based reweighting analysis method) algorithm,^20^ as implemented in the PyEMMA software package,^40^ was used to construct multi-ensemble Markov Models of apoMb from the simulation trajectory data. TRAM uses information from a series of thermodynamic ensembles, labeled by index *k*, to infer both the transition rates 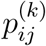 between states *i* and *j* in ensemble *k* and the conformational (reduced) free energies 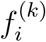 of each state *i* in ensemble *k*. This is achieved by maximization of a joint likelihood function *L*_TRAM_ that is the product of a reversible MSM estimator likelihood function 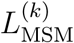, and a free energy estimator likelihood function 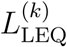:

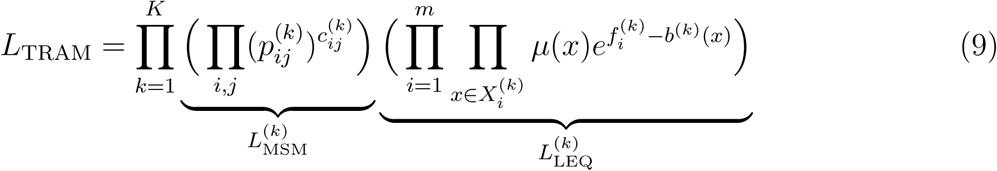

where 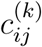 are the number of transition counts between states *i* and *j* observed in ensemble *k, µ*(*x*) is the normalized equilibrium probability (Σ*_x_ µ*(*x*) = 1) of each sample *x*, 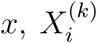 is the set of samples *x* drawn from the *k*^th^ ensemble, and *b*^(*k*)^(*x*) is the (reduced) bias potential acting on sample *x* in ensemble *k*.

Due to insufficient overlap of sampled conformational states across the simulated temperatures, MEMMs were constructed for each temperature, using all simulation trajectory data obtained at all viable restraint biases. For example, for simulations performed at 350 K, a total of four restraint-biased ensembles were included in the TRAM estimation, for force constant values of *k* = 0.5, 0.7, 1.0, and 1.2 kJ. The key quantity of interest resulting from these calculations are the equilibrium populations of conformational states.

We projected the 350 K trajectory data from each microstate onto tIC_1_ after weighting by its estimated population, to obtain a series of free energy profiles *F* ^(*k*)^(tIC_1_) = *-k*_*B*_*T* ln *π*^(*k*)^(tIC_1_) for each thermodynamic ensemble *k*. All free energy profiles show two minima separated by a 2–4 kcal mol^*-*1^ barrier, with the global minimum shifting from structured conformations to unstructured conformations for helix F as the force constant increases (Figure 9). This clearly shows how the restraint biases are able to achieve a range of conformational distributions which we can reconcile against experimental measurements.

**Figure 9:**
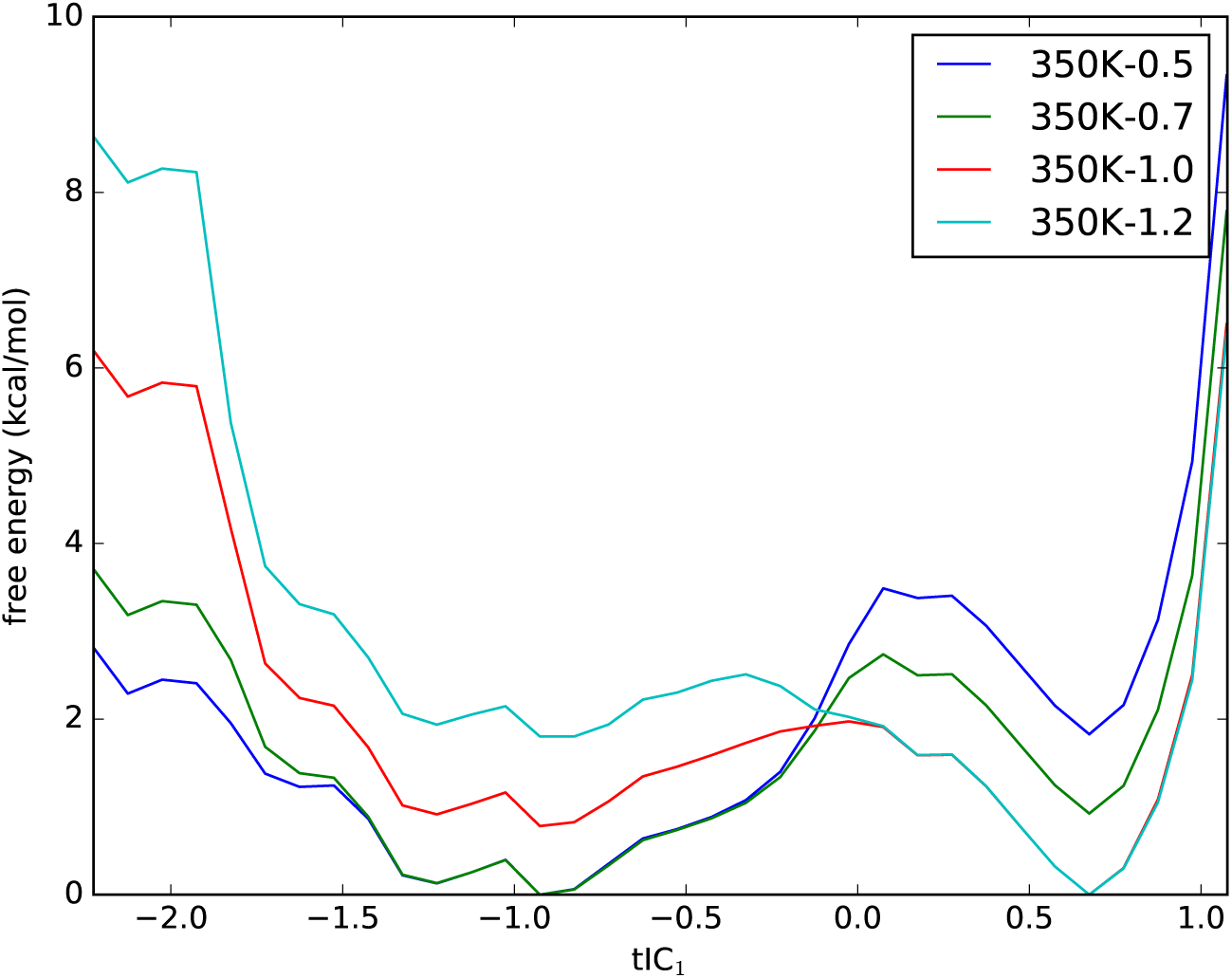
TRAM predictions of free energy profiles along tIC_1_ for ensembles at 350 K and with biases of *k* = 0.5, 0.7, 1.0, and 1.2 kJ.

### Part III: Reconciling multi-ensemble MSMs against experimental HDX protection factor and chemical shifts data using BICePs

From the work described in the previous section (Part II), we now have in hand a series of MSMs for each thermodynamic ensemble (defined by a particular temperature and restraint bias), each yielding a prediction of equilibrium conformational populations. From the work described in Part I, we also have in hand a predictor of the observable (ln PF_*i*_) for each conformational state *X*,

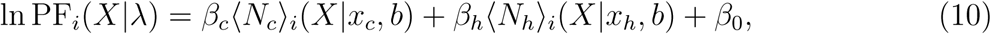

along with the full posterior *P* (*λ*) of nuisance parameters *λ* = (*β*_*c*_, *β*_*h*_, *β*_0_, *x*_*c*_, *x*_*h*_, *b*).

Using these two ingredients, we now proceed to reconcile each MSM against the experimentally measured observables. Specifically, we wish to find the set of MSM-predicted conformational populations that best agrees with the experimental observables. To do accomplish this task, we take a Bayesian inference approach, implemented through an algorithm we call Bayesian Inference of Conformational Populations, or BICePs.

#### The BICePs algorithm

For a full discussion of the background and development of BICePs, please refer to our previous work.^21,22,41,42^

The purpose of BICePs is to make unbiased estimates of conformational populations by optimally combining information from theoretical predictions (here, all-atom simulations) and ensemble-averaged experimental observables. The goal is to sample the posterior probability distribution of conformational states *X*, given some experimental data *D*. By Bayes’ Theorem,

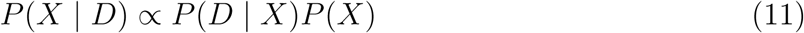

where *P* (*D* | *X*) is a likelihood function representing experimental restraints, and *P* (*X*) is a prior probability function, calculated from a theoretical model (in this case, from molecular simulation). BICePs is similar to other Bayesian methods for the inference of structural ensembles, including ISD,^43^ MELD,^44^ and Metainference. ^45^

#### Nuisance parameters

One important feature of BICePs is the ability to infer how best to balance the relative influence of experimental versus theoretical restraints. It does this by modeling the (unknown) uncertainty of the experimental measurements and heterogeneity in the conformational ensemble using nuisance parameters *σ*, and sampling over these parameters to estimate their posterior distribution as well, through

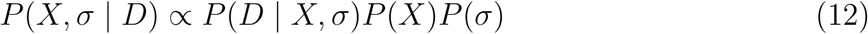

where we assume some prior distribution *P* (*σ*).

#### Reference potentials

Another important feature of BICePs is the use of reference potentials. An experimental observable *r*(*X*) is a projection of a high-dimensional conformational ensemble *X* to a single-valued function, and therefore the likelihood *P* (*D* | *X, σ*) of observing a particular value *r*(*X*) must be expressed relative to some reference probability *P*_ref_(*r*(*X*)) of observing *r*(*X*) in the absence of any particular structure, according to

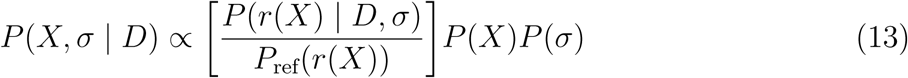

#### BICePs scores for unbiased model selection

As discussed by Ge et al.,^22^ another advantage of BICePs is its ability to perform model selection. Given a set of conformational populations predicted by an MSM, we wish to objectively evaluate the extent to which it agrees with experimental observables, and be able to rank it against other models.

Suppose we are presented with a collection of competing models *P* ^(*k*)^(*X, σ* | *D*), each with a different theoretical prior *P* ^(*k*)^(*X*) predicted from an MSM. The total evidence for model *P* ^(*k*)^ can be expressed as

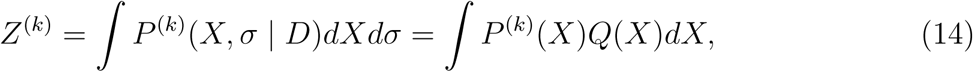

where *Q*(*X*) = ∫ [*P* (*r*(*X*) | *D, σ*)*/P*_ref_(*r*(*X*))]*P* (*σ*)*dσ* represents the probability of *X* given the experimental data. As can be seen by the last term in Equation (14), *Z*^(*k*)^ is an overlap integral that quantifies how well the theoretical *P* ^(*k*)^(*X*) agrees with the experimental data.

To compare two different models *P* ^(1)^ and *P* ^(2)^, it is common to compute the ratio of total evidences, *Z*^(1)^*/Z*^(2)^, often called the *Bayes factor*. To facilitate the assignment of a unique score to each model, we compute a Bayes factor where the second model is a “null” model *Z*_0_ in which *P* ^(*k*)^(*X*) is a uniform distribution of conformational states. In this way, we define a quantity we call the BICePs score, *f* ^(*k*)^, for each model *P* ^(*k*)^,

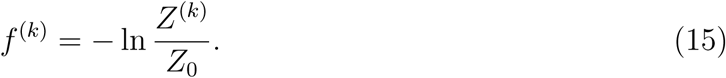

In practice, the calculation of the BICePs score *f* ^(*k*)^ can be performed using free energy estimation techniques, and can be thought of as a “free energy” of each model *P* ^(*k*)^; the lower the value of *f* ^(*k*)^, the better the model agrees with the experimental data. We can thus use the BICePs score *f* ^(*k*)^ for objective model selection. We use the multistate Bennett acceptance ratio (MBAR) method^46^ was used to calculate the BICePs scores *f* ^(*k*)^.

### Reconciling conformational populations of apoMb against experimental observables using BICePs

Here, we use two kinds of experimental data with the BICePs algorithm: HDX protection factors measured by Nishimura et al., ^37^ and NMR chemical shifts for H, C_*α*_, and N atoms measured by Eliezer and Wright.^30^ Experimental protection factor data for apoMb, converted to ln PF values, are listed in the Supporting Information (Table S3).

#### Protection factor restraints

For each residue *i*, we introduce a Gaussian function to restrain the computed observable *r*_*i*_(*X*) = ln PF_*i*_(*X*), against the measured values 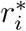,

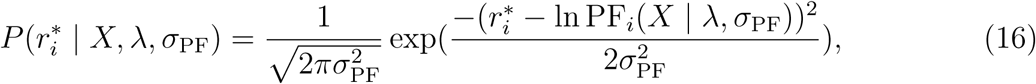

where *σ*_PF_ represents the uncertainty in the experimental measurement, and *λ* = (*β*_*c*_, *β*_*h*_, *β*_0_, *x*_*c*_, *x*_*h*_, *b*). As in previous BICePs calculations, ^21,42^ we used exponential reference potentials *P*_ref_(*r*_*i*_(*X*)) for all residues *i*, and an uninformative Jeffreys prior 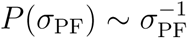. The prior distribution *P* (*λ*) comes from the posterior distribution of the nuisance parameters sampled in Part I (see Figure 2).

#### Chemical shift restraints

For each residue *i*, we introduce a Gaussian function to restrain predicted chemical shift values *δ*_*i*_(*X*) for each conformational state *X*, against the measured values 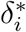,

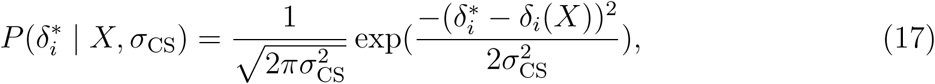

where *σ*_CS_ represents the uncertainty in the chemical shift measurement. Predictions of NMR chemical shifts *δ*_*i*_(*X*) for each conformational state *X* were calculated using the SHIFTX2 algorithm^47^ as implemented in MDTraj,^48^ using the default user-specified parameters of pH 7.0 and 298 K. The predicted chemical shifts *δ*_*i*_(*X*) are the ensemble-averaged values of predictions of each trajectory snapshot belonging to state *X*. Exponential reference potentials *P*_ref_(*δ*_*i*_(*X*)) were used for all residues *i*, along with an uninformative Jeffreys prior for *P* (*σ*_CS_).

#### Sampling the posterior distribution with BICePs

Taking together the protection factor and chemical shift data, the full posterior function is proportional to the product a prior *P* (*X*) (i.e. the predicted conformational state populations), all four likelihood functions, and priors for all nuisance parameters:

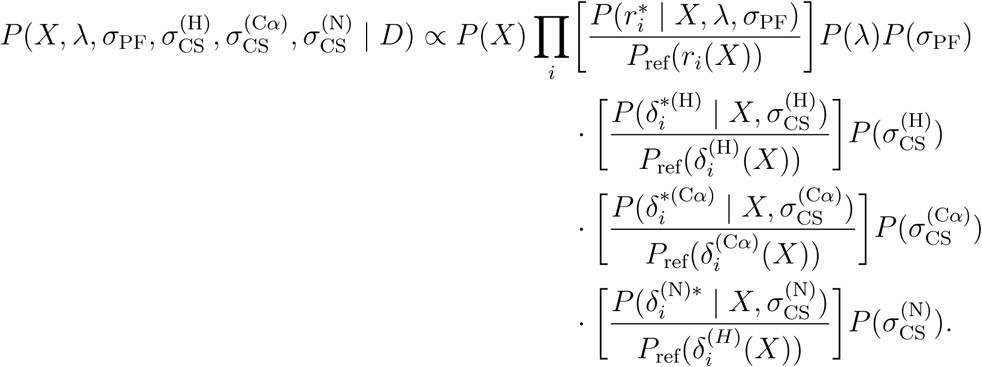

To sample this posterior probability function, 10^7^ steps of Markov Chain Monte Carlo (MCMC) was performed. We employed several strategies to make this sampling efficient. First, a finite grid of possible values of 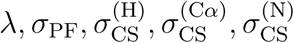 were used, where the sums of squared errors (SSE) were precomputed. Proposed MCMC moves were allowed to neighboring values in the grid. Grid values for parameters with Jeffreys priors were log-scaled to enforce the prior and improve acceptance ratios.

The number of grid values for each nuisance parameter was chosen to keep the acceptance ratio around 0.5. The dimensions of the array storing SSE values for *λ* = *β*_*c*_, *β*_*h*_, *β*_0_, *x*_*c*_, *x*_*h*_, *b* was carried over from Part I. Although large (20 × 26 × 50 × 7 × 8 × 18 = 26.2 million values), the array could be stored in memory.

### BICePs scores quantify the conformational ensembles that best agree with experimental data

As mentioned above, quantitative comparison between different models can be performed using the BICePs score. From the work in Part II, we have 31 models of the prior population distribution *P* ^(*k*)^(*X*) calculated using TRAM, each at different temperatures and different restraint biases. For each of these, a BICePs calculation was performed to sample the posterior distribution *P* ^(*k*)^(*X*|*D*), where *D* is the experimental data, and the BICePs score was computed to rank the model.

To evaluate the effects of including chemical shift data, these calculations were repeated two-fold: once using only the protection factor (PF) experimental data, and once using both the protection factor and chemical shift data (PF+CS) (Figure 10). The results show that for each simulated temperature, there is a restraint bias for which the sampled conformational ensemble achieves the best overlap with experimental restraints. In all cases, BICePs scores calculated using PF+CS data are lower than for those calculated using PF data alone. This indicates that the additional use of CS data yields models that agree better with experiment than PF data alone.

**Figure 10:**
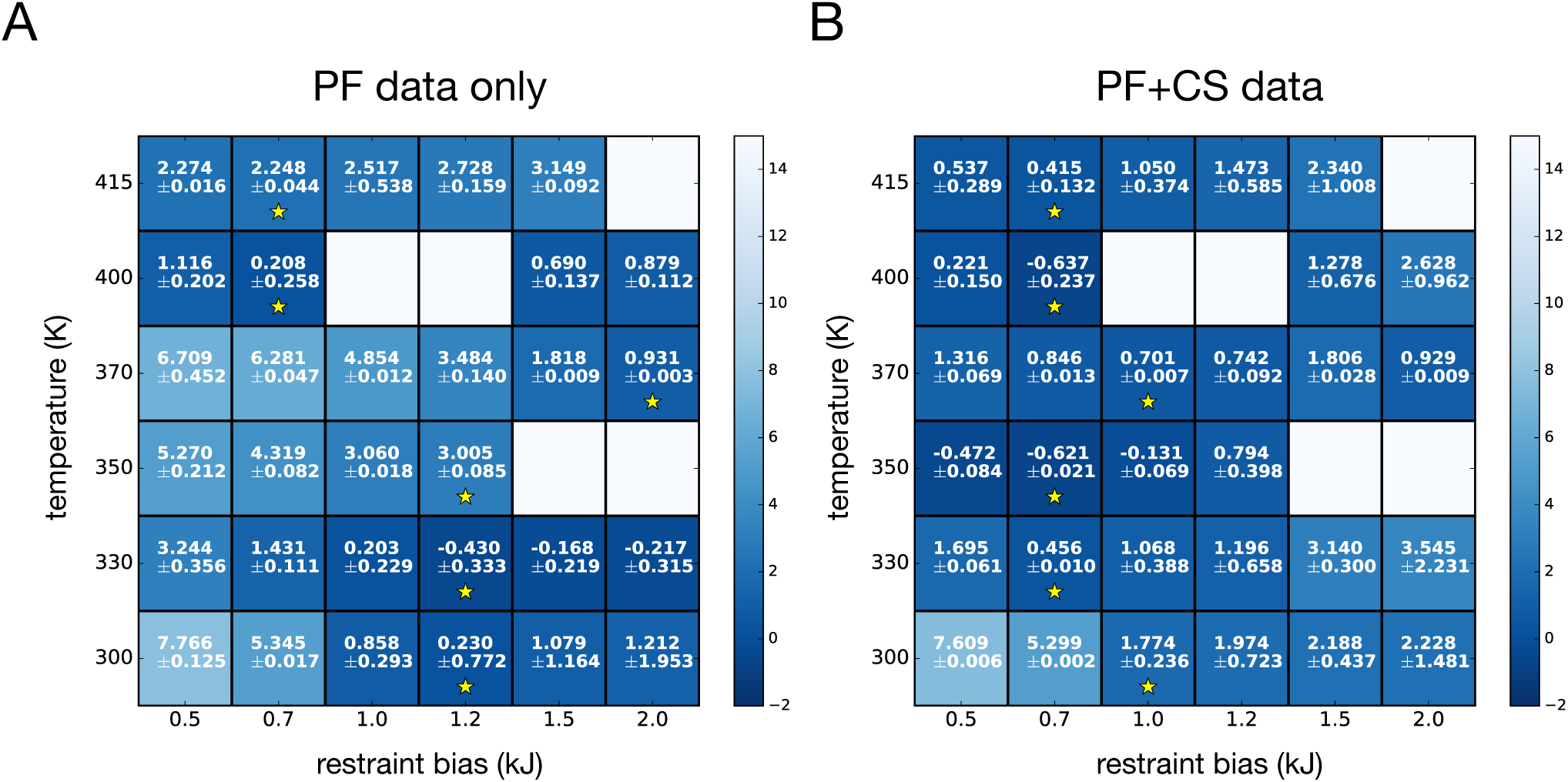
A summary of the BICePs scores computed for MEMMs built at different temperatures and restraint bias potentials, for BICePs calculations performed using (A) only protection factor data (PF), and (B) protection factor and chemical shift data (PF+CS). Each cell shows the computed BICePs score, with uncertainty estimates computed as standard deviations over 5 rounds of 10^7^-step MCMC sampling. Blank cells indicate no models for those restraint biases were constructed due to the lack of viable trajectory input for the TRAM calculation. Cells with yellow stars mark the best model at each temperature.

When we examine the restraint bias corresponding to the best model for each temperature (Figure 10, yellow stars), we find differences between the PF and PF+CS calculations. For the PF BICePs calculations, a majority of the best models correspond to restraint bias of 1.2 kJ, while for the PF+CS calculations, a majority correspond to a restraint bias of 0.7 kJ. We believe this may be because a gentler restraint bias is needed to produce conformations with intact secondary structure in better agreement with measured chemical shift data.

A comparison of the best models at each temperature is shown in Figure 11. The model with the lowest BICePs score is one corresponding to the 400 K simulations, using a restraint bias of 0.7 kJ. For this model, we compared the populations predicted by BICePs using only the experimental data (i.e. a uniform prior *P* ^(*k*)^(*X*)) against BICePs predictions using both the experimental data and the prior given by the TRAM calculation (Figure 12). We find that, in each scenario, conformational state 18 has the dominant population. This state has an intermediate extent of structure in helix F and H, located near the middle of the tICA landscape (see Figure 8) The second-highest population, conformational state 21, has more disorder in helix F and H and is located near the right side of the tICA landscape. Using experimental restraints alone, the population of state 21 is estimated at around 15%; the prior given by the TRAM calculation, however, increases the predicted posterior population to more than 30% (Figure 12b). Other states are predicted to contribute much less population (under 1%).

**Figure 11:**
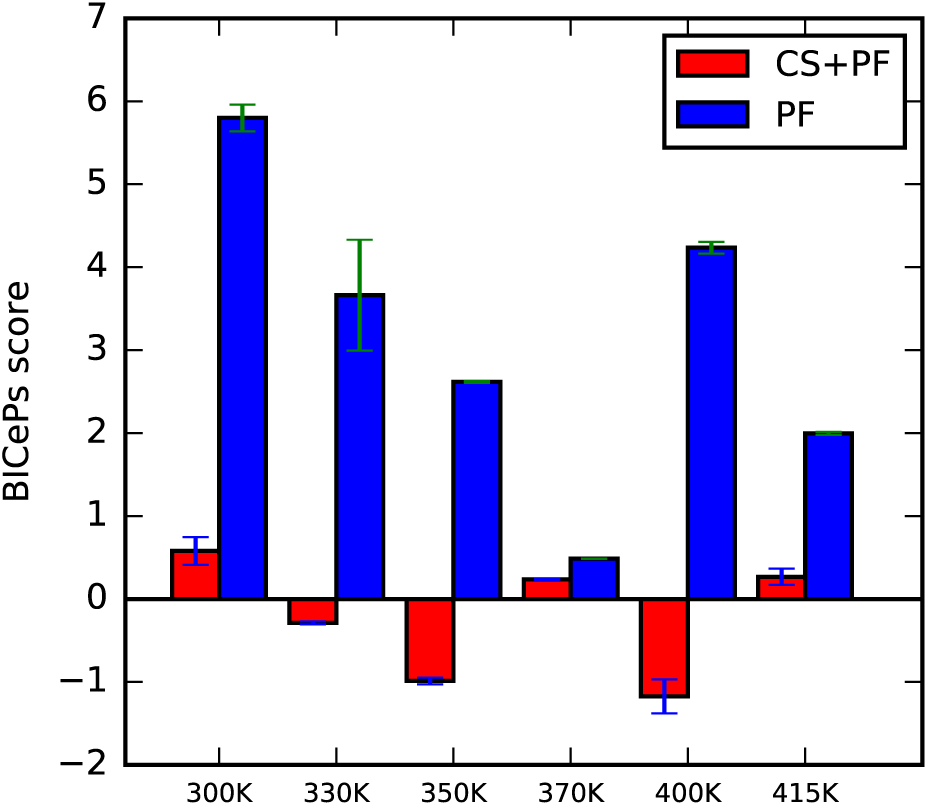
A comparison of the best BICePs scores at each temperature, for BICePs calculations performed with only PF restraints (blue) and PF+CS restraints (red). Error bars are standard deviations of BICePs scores computed over 5 rounds of 10^7^-step MCMC sampling.

**Figure 12:**
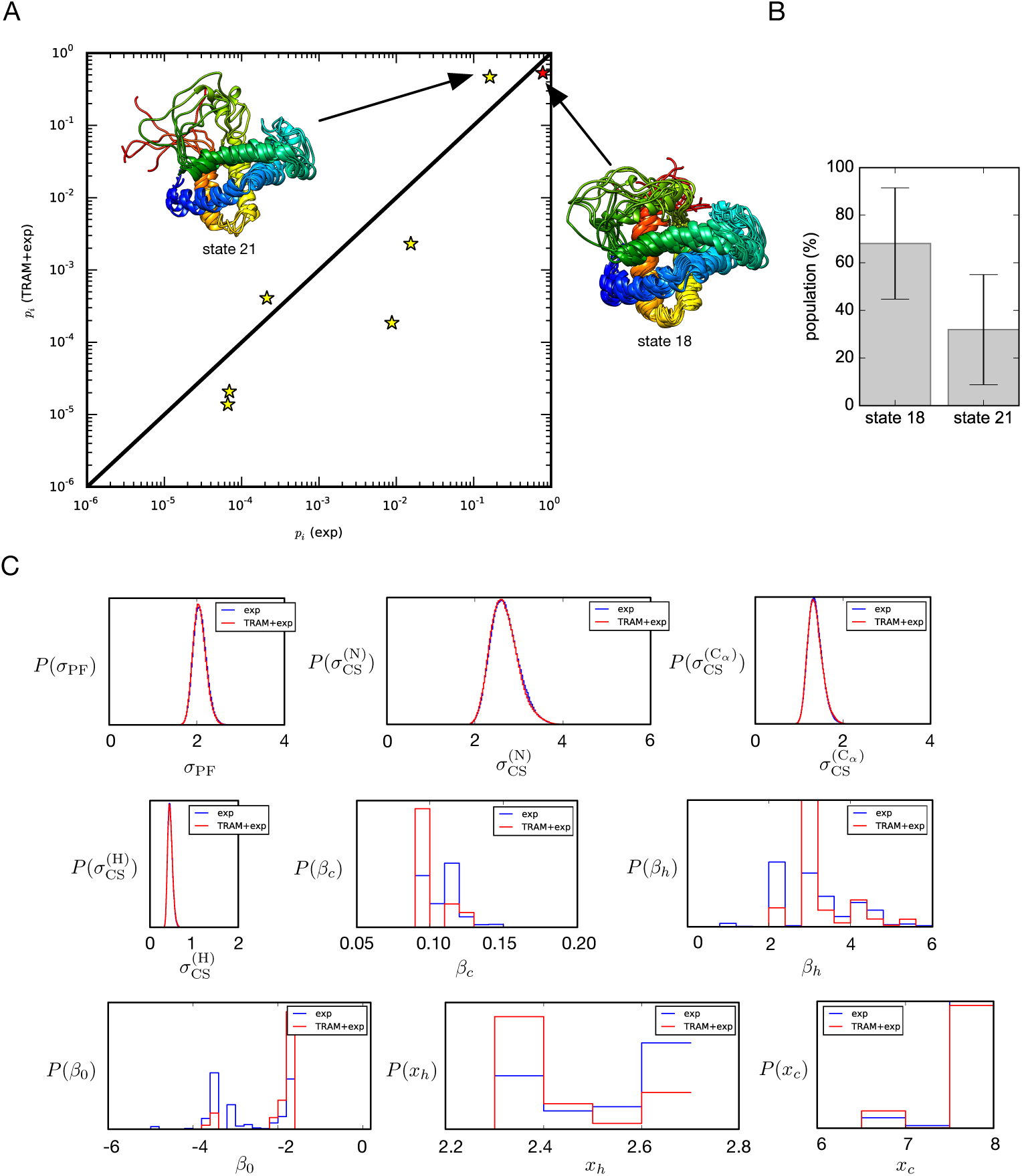
Posterior distributions of conformational populations and model parameters for the best-scoring model (400 K, 0.7 kJ). (A) A comparison of posterior conformational state populations *p*_*i*_ predicted by BICePs in the absence of simulation information (exp, i.e. a uniform prior of state populations), versus populations predicted by BICePs using both experimental information and prior conformational state populations from the best-scoring model (TRAM+exp). (B) Estimated populations of the two dominant conformational states in the model. Error bars are standard deviations of BICePs scores over 5 rounds of 10^7^-step MCMC sampling. (C) Posterior distributions of all nuisance parameters sampled by BICePs, for both exp and TRAM+exp scenarios.

The predicted state populations (Figure 12a) can be thought of as the marginal posterior distribution *P* (*X*) sampled by BICePs. Marginal posterior distributions of nuisance parameters sampled by BICePs are shown in Figure 12c. For the 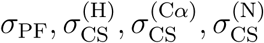 parameters, the posterior distributions give estimates of the standard errors in comparing the experimental observables against the predictions. For example, the standard error when comparing the ln PF_*i*_ values against the experimental values is about two natural logarithm units. The standard error when comparing experimental versus computed chemical shifts is about 0.5, 1.1, and 2.6 ppm for H, C_*α*_ and N chemical shifts, respectively. The sampled posterior distributions for the 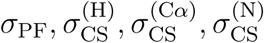 parameters are nearly identical in the two cases when only experimental restraints included (exp), and when both simulation and experimental data is included (TRAM+exp). The sampled posterior distributions for the other nuisance parameters *λ* = (*β*_*c*_, *β*_*h*_, *β*_0_, *x*_*c*_, *x*_*h*_, *b*) closely follow the posterior distributions sampled in Part I.

Using the mixture of (TRAM+exp) populations sampled by BICePs, we calculated ensemble-averaged predictions of protection factor and chemical shift observables, enabling direct comparison to the experimental values as a function of residue index (Figure 13). The results show excellent agreement with all experimental observables. The experimental and simulated N, C_*α*_ and H chemical shifts have squared correlation coefficients of *R*^2^ = 0.84, and 0.66, respectively. The good agreement is somewhat expected, as the chemical shift deviations largely report secondary structure in the structured regions of helices A, B, C, D, E and G, which is largely intact in our simulations. More remarkable is the agreement between simulated and experimental protection factors: the squared correlation coefficient for experimental and simulated values of ln PF_*i*_ is *R*^2^= 0.72, which rivals Persson and Halles results for BPTI (*R*^2^ = 0.68).^17^

**Figure 13:**
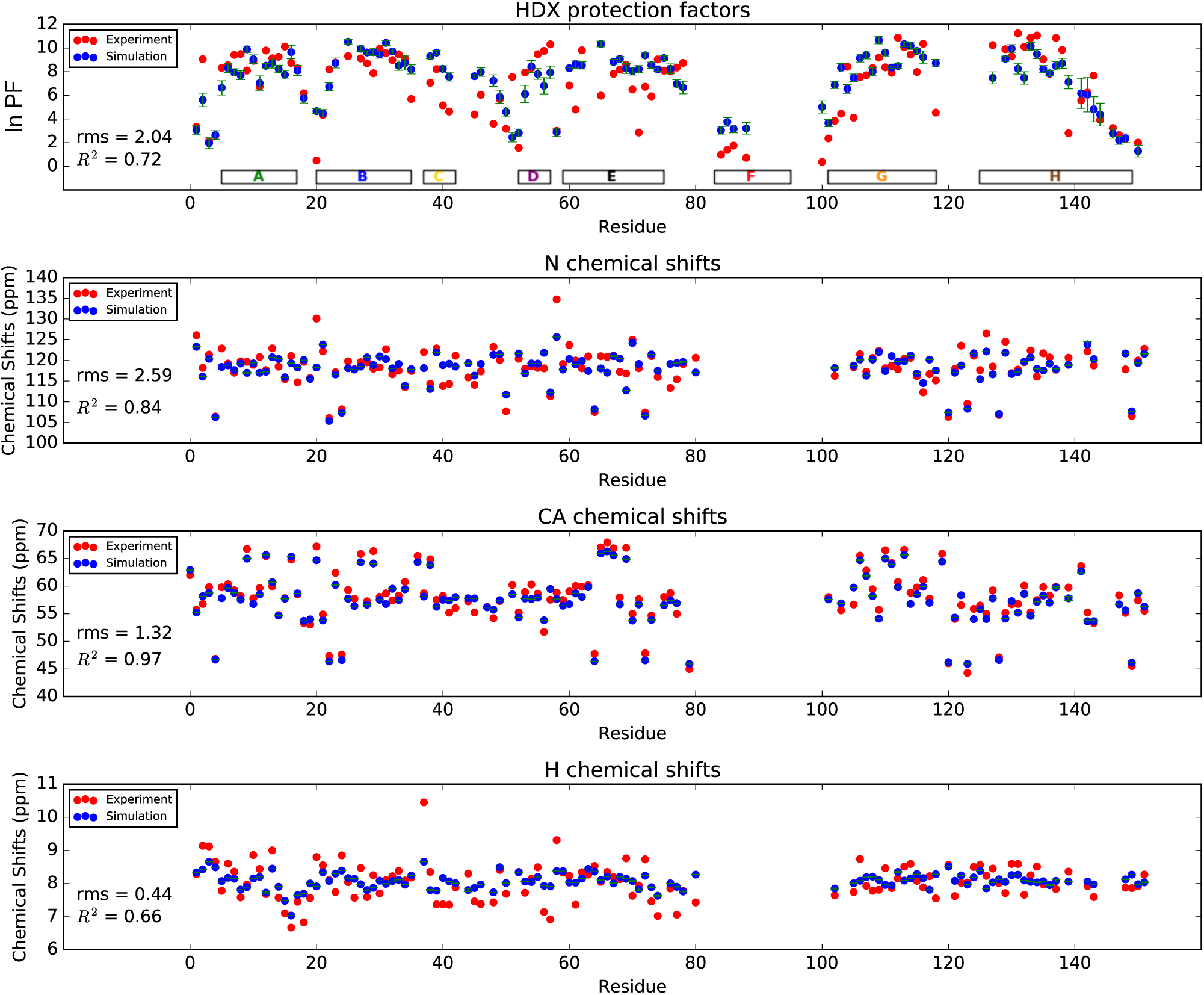
Residue profiles of experimental and simulated protection factor and chemical shift observables for apomylobin, using the conformational state populations of the bestscoring BICePs model (400 K, 0.7 kJ).

## Discussion

Using the methodologies described in Parts I, II and III, we have constructed a number of models of the apomyoglobin native state with different populations of conformational states, and used Bayesian inference to interrogate how well each model predicts experimental HDX protection factor and chemical shift observables. Our best model is dominated by a 70%-30% mixture of two conformational states (18 and 21), the first of which has a partially disordered yet compact helix F, and the second of which has a more disordered and solvent-exposed helix F.

Experimentally, chemical shifts are not reported for helix F residues in the apoMb Nstate (pH 6.1).^30^ This is because of the slow chemical exchange of these residues, presumably due to fluctuations between conformational substates. Our model, which describes multiple populations of heterogeneous conformational states, is consistent with this picture.

While here we are focused on using the TRAM method to predict equilibrium populations, this method can also be used to make predictions for the kinetics of interconversion between states. Thus, the methods developed in this work offer a way to get information about conformational dynamics from thermodynamically averaged experimental observables.

Another way to infer kinetic information from models of conformational state populations would be to (1) construct Markov model of dynamics in the absence of experimental information, (2) use BICePs (or some other method) to estimate improved state populations given ensemble-averaged experimental observables, and finally (3) use Maximum-Caliber method to infer changes in the transition rates between the conformational states. ^49^ These methods could complement and improve existing methods such as augmented Markov models ^50^ by providing a fully Bayesian approach to inferring model parameters.

## Conclusion

In this work we have presented, in three parts, new and improved ways of reconciling simulated ensembles of protein conformations against experimental observables and applied them to modeling the N-state of apomyoglobin using HDX protection factor measurements. First, we have parameterized a new empirical predictor of HDX protection factors based on structural observables from simulation trajectory data, and applied Bayesian inference to infer the complete posterior distribution of nuisance parameters. Importantly, we show that the posterior probability gives improved results, distinct from a simple “best-fit” model.

We have also presented a new way to use bias potentials in molecular simulations to sample solvent-exposed conformations. We use this method to construct a series of multiensemble Markov State Models of apomyoglobin, resulting in a number of candidate models consisting of metastable conformational states and their populations.

Finally, we have used the BICePs algorithm to reconcile each model against experimental protection factor and chemical shift observables, using BICePs scores to objectively select the best model of the apomyoglobin N state. The best-scoring model is dominated by two conformational substates: one with partially disordered and compact helix F, and another with a more disordered and solvent-exposed helix F. This model agrees well with experimental protection factors (*R*^2^ = 0.72), and is consistent with the observation of slow chemical exchange in the helix F region.

These tools offer new ways to refine conformational ensembles against protection factor data, utilizing the framework of Bayesian inference.

## Acknowledgement

The authors thank Heinrich Roder for valuable expertise and advice, Juliette Lacomte for NMR structure refinement information for sperm whale apomyoglobin at pH 7, and D. E. Shaw Research for providing access to folding trajectory data. This work was supported by the National Science Foundation grant number MCB-140270, and in part through major research instrumentation grant number CNS-09-58854. Computing resources were supported through the Extreme Science and Engineering Discovery Environment (XSEDE) Stampede supercomputer at the Texas Advanced Computing Center (TACC).

## Supporting Information Available

Supporting Tables S1, S2 and S3.

## Supporting Information

### Supporting Tables

**Table S1:** Experimental protection factors measured for ubiquitin taken from Craig et al., ^23^ converted to ln PF values.

| residue | number | $\ln$ PF |
| --- | --- | --- |
| GLN | 2 | 6.210072 |
| ILE | 3 | 13.7372227 |
| PHE | 4 | 13.4839383 |
| VAL | 5 | 13.1523661 |
| LYS | 6 | 10.6909026 |
| THR | 7 | 10.4629467 |
| LEU | 8 | 0.67005226 |
| THR | 9 | 0 |
| GLY | 10 | 3.93511792 |
| LYS | 11 | 5.21075007 |
| THR | 12 | 3.2443424 |
| ILE | 13 | 11.0570136 |
| THR | 14 | 2.53514619 |
| LEU | 15 | 11.5336487 |
| GLU | 16 | 5.14167251 |
| VAL | 17 | 12.2405424 |
| GLU | 18 | 7.97845735 |
| SER | 20 | 3.82459384 |
| ASP | 21 | 12.9796722 |
| THR | 22 | 7.73438333 |

**Table S1 – continued from previous page**
| residue | number | ln PF |
| --- | --- | --- |
| ILE | 23 | 11.2135894 |
| GLU | 24 | 5.00581999 |
| ASN | 25 | 10.5550501 |
| VAL | 26 | 14.6214153 |
| LYS | 27 | 14.8539764 |
| ALA | 28 | 11.4806893 |
| LYS | 29 | 13.4517021 |
| ILE | 30 | 14.2483966 |
| GLN | 31 | 7.7689221 |
| ASP | 32 | 5.6229128 |
| LYS | 33 | 4.56602624 |
| GLU | 34 | 6.21467717 |
| GLY | 35 | 6.26763662 |
| ILE | 36 | 6.981438 |
| ASP | 39 | 1.64174317 |
| GLN | 40 | 6.45875119 |
| GLN | 41 | 8.26167531 |
| ARG | 42 | 9.13435506 |
| LEU | 43 | 6.56927527 |
| ILE | 44 | 12.6573103 |
| PHE | 45 | 6.49328996 |
| ALA | 46 | 0 |
| GLY | 47 | 4.1699816 |
| LYS | 48 | 7.86102551 |

**Table S1 – continued from previous page**
| residue | number | ln PF |
| --- | --- | --- |
| GLN | 49 | 3.80617316 |
| LEU | 50 | 8.45969763 |
| GLU | 51 | 5.41798272 |
| ASP | 52 | 2.73316851 |
| GLY | 53 | 3.31802512 |
| ARG | 54 | 9.9932193 |
| THR | 55 | 11.3494419 |
| LEU | 56 | 13.0211187 |
| SER | 57 | 6.68440452 |
| ASP | 58 | 6.84098031 |
| TYR | 59 | 10.9580025 |
| ASN | 60 | 5.11404149 |
| ILE | 61 | 8.54949845 |
| GLN | 62 | 8.27088565 |
| LYS | 63 | 3.09927954 |
| GLU | 64 | 6.37125295 |
| SER | 65 | 7.52484808 |
| THR | 66 | 6.1939539 |
| LEU | 67 | 7.75740918 |
| HIS | 68 | 8.49884158 |
| LEU | 69 | 7.95773408 |
| VAL | 70 | 9.09981629 |
| LEU | 71 | 3.0278994 |
| ARG | 72 | 2.5788953 |

**Table S1 – continued from previous page**
| residue | number | ln PF |
| --- | --- | --- |
| LEU | 73 | 0 |
| ARG | 74 | 0 |
| GLY | 75 | 0 |
| GLY | 76 | 0 |

**Table S2:** Experimental protection factors measured for BPTI taken from Persson et al., ^17^ converted to ln PF values.

| residue | number | ln PF |
| --- | --- | --- |
| CYS | 5 | 8.52877518 |
| LEU | 6 | 7.43504727 |
| GLU | 7 | 8.229439 |
| TYR | 10 | 5.756463 |
| GLY | 12 | 3.840712 |
| ALA | 16 | 6.963017 |
| ARG | 17 | 1.752267 |
| IIE | 18 | 12.37639 |
| IIE | 19 | 2.256533 |
| ALA | 25 | 3.04632 |
| GLY | 28 | 7.676819 |
| LEU | 29 | 10.85899 |
| CYS | 30 | 3.677228 |
| THR | 32 | 5.701201 |
| VAL | 34 | 3.734793 |
| TYR | 35 | 11.23431 |

**Table S2 – continued from previous page**
| residue | number | ln PF |
| --- | --- | --- |
| GLY | 36 | 9.574149 |
| GLY | 37 | 11.43924 |
| CYS | 38 | 4.503856 |
| LYS | 41 | 6.988346 |
| ARG | 42 | 2.141404 |
| ASN | 43 | 5.125554 |
| ASN | 44 | 14.02274 |
| SER | 47 | 4.503856 |
| ALA | 48 | 2.403899 |
| MET | 52 | 11.02938 |
| ARG | 53 | 9.825131 |
| THR | 54 | 7.962339 |
| CYS | 55 | 12.1139 |
| GLY | 56 | 8.008391 |

**Table S3:** Experimental protection factors measured for apomyoglobin at pH 6, taken from Nishimura et al., ^37^ and converted to ln PF values.

| residue | number | ln PF |
| --- | --- | --- |
| LEU | 2 | 3.3387 |
| SER | 3 | 9.0552 |
| GLU | 4 | 2.1162 |
| GLY | 5 | 2.6101 |
| GLU | 6 | 8.3181 |
| TRP | 7 | 8.5457 |

**Table S3 – continued from previous page**
| residue | number | ln PF |
| --- | --- | --- |
| GLN | 8 | 9.4170 |
| LEU | 9 | 9.4915 |
| VAL | 10 | 8.0990 |
| LEU | 11 | 8.9169 |
| HIS | 12 | 6.7276 |
| VAL | 13 | 9.7889 |
| TRP | 14 | 9.1266 |
| ALA | 15 | 9.2542 |
| LYS | 16 | 10.1224 |
| VAL | 17 | 8.7396 |
| GLU | 18 | 8.2774 |
| ALA | 19 | 6.1741 |
| VAL | 21 | 0.5004 |
| ALA | 22 | 4.3512 |
| GLY | 23 | 8.1983 |
| HIS | 24 | 8.7599 |
| GLN | 26 | 9.3119 |
| ILE | 28 | 9.1152 |
| LEU | 29 | 8.6984 |
| ILE | 30 | 7.8670 |
| ARG | 31 | 9.9290 |
| LEU | 32 | 9.5716 |
| PHE | 33 | 8.9773 |
| LYS | 34 | 9.4717 |

**Table S3 – continued from previous page**
| residue | number | ln PF |
| --- | --- | --- |
| SER | 35 | 9.0890 |
| HIS | 36 | 5.6884 |
| THR | 39 | 7.0556 |
| LEU | 40 | 8.2188 |
| GLU | 41 | 5.1578 |
| LYS | 42 | 4.6338 |
| PHE | 46 | 4.3853 |
| LYS | 47 | 6.0469 |
| LEU | 49 | 3.6004 |
| LYS | 50 | 5.6584 |
| THR | 51 | 3.1733 |
| GLU | 52 | 7.5530 |
| ALA | 53 | 1.5526 |
| GLU | 54 | 7.9263 |
| MET | 55 | 8.3330 |
| LYS | 56 | 9.4799 |
| ALA | 57 | 9.7606 |
| SER | 58 | 10.3153 |
| GLU | 59 | 3.0033 |
| LEU | 61 | 6.8314 |
| LYS | 62 | 4.8015 |
| LYS | 63 | 9.8059 |
| VAL | 66 | 5.9719 |
| VAL | 68 | 7.8175 |

**Table S3 – continued from previous page**
| residue | number | ln PF |
| --- | --- | --- |
| LEU | 69 | 8.1635 |
| THR | 70 | 8.5773 |
| ALA | 71 | 6.4889 |
| LEU | 72 | 2.8538 |
| GLY | 73 | 6.7213 |
| ALA | 74 | 5.9142 |
| ILE | 75 | 9.0367 |
| LEU | 76 | 8.0870 |
| LYS | 77 | 8.0141 |
| LYS | 78 | 8.3490 |
| LYS | 79 | 8.7400 |
| GLU | 85 | 0.9784 |
| LEU | 86 | 1.3868 |
| LYS | 87 | 1.7472 |
| LEU | 89 | 0.7190 |
| ILE | 101 | 0.3748 |
| LYS | 102 | 2.3550 |
| TYR | 103 | 3.8462 |
| LEU | 104 | 4.4498 |
| GLU | 105 | 8.4099 |
| PHE | 106 | 4.1129 |
| ILE | 107 | 7.5292 |
| SER | 108 | 7.6937 |
| GLU | 109 | 8.3079 |

**Table S3 – continued from previous page**
| residue | number | ln PF |
| --- | --- | --- |
| ALA | 110 | 9.1859 |
| ILE | 111 | 8.3689 |
| ILE | 112 | 7.9042 |
| HIS | 113 | 10.8719 |
| VAL | 114 | 10.0759 |
| LEU | 115 | 9.4460 |
| HIS | 116 | 7.9725 |
| SER | 117 | 10.3608 |
| HIS | 119 | 4.5392 |
| GLN | 128 | 10.2523 |
| ALA | 130 | 9.8983 |
| MET | 131 | 9.2917 |
| ASN | 132 | 11.2423 |
| LYS | 133 | 10.1108 |
| ALA | 134 | 10.8549 |
| LEU | 135 | 11.0577 |
| GLU | 136 | 9.0395 |
| LEU | 137 | 7.8229 |
| PHE | 138 | 10.8617 |
| ARG | 139 | 9.8380 |
| LYS | 140 | 2.7987 |
| ILE | 142 | 5.5787 |
| ALA | 143 | 6.2290 |
| ALA | 144 | 7.6644 |

**Table S3 – continued from previous page**
| residue | number | ln PF |
| --- | --- | --- |
| LYS | 145 | 3.9210 |
| LYS | 147 | 3.2341 |
| GLU | 148 | 2.6429 |
| LEU | 149 | 2.3232 |
| TYR | 151 | 2.0006 |

